# Genome assembly variation and its implications for gene discovery in nematode species

**DOI:** 10.1101/2024.02.26.582167

**Authors:** Grace M. Mariene, James D. Wasmuth

## Abstract

Genome assemblers are a critical component of genome science, but the choice of assembly software and protocols can be daunting. Here, we investigate genome assembly variation and its implications for gene discovery across three nematode species—*Caenorhabditis bovis*, *Haemonchus contortus*, and *Heligmosomoides bakeri*—highlighting the critical interplay between assembly choice and downstream genomic analysis. Selecting popular genome assemblers, we generated multiple assemblies for each species, analyzing their structure, completeness, and effect on gene family analysis. Our findings demonstrate that assembly variations can significantly affect gene family composition, with notable differences in critical gene families like *cyp*, *<u>gst</u>*, *ugt*, and *nhr*. Despite broadly similar performance using various assembly metrics, comparisons of assemblies with a single species revealed underlying structural rearrangements and inconsistencies in gene content. This emphasizes the imperative for continuous refinement of genomic resources. Our findings advocate for a cautious and informed approach to genome assembly and annotation to ensure reliable and insightful genomic interpretations.

## Introduction

Genome assembly software serves a critical, often underappreciated role in the life sciences. Its purpose is to reconstruct, *de novo*, the complete sequence of an organism’s DNA—its genome—from shorter DNA fragments obtained through one or more sequencing technologies. A typical assembler performs various tasks, such as error correction, read alignment and overlap detection, and consensus building. The decrease cost of sequencing has empowered the launch of large BioGenome consortia which aim to sequence, assemble and annotation the genomes of as many animal and plant species as possible (Mieszkowska et al., 2022; Ebenezer et al., 2022). Individual research groups are also able to sequence the genomes of their favourite species. This places the evolution and refinement of genome assemblers at the core of current and future breakthroughs in health, agricultural, and environmental sciences.

Understanding accuracy of the output of the genome assembler—long strings of As, Gs, Cs, Ts and the occasional N—is helpful, as errors in the produced assembly will carry into and magnify through the annotation and analysis, often without notice. There are many sources of error. One example is high and/or inconsistent levels of heterozygosity in the organism’s genome leading to sequence reads being assembled as separate loci rather than collapsed in the haploid assembly. A second example is long regions of repetitive DNA and transposable elements scatered through the genome being incorrectly collapsed or rearranged, leading to gaps. While some genome assemblers specialise in one particular problem, for example heterozygosity (Kajitani et al., 2014), most assemblers are generalists and seek to identify and resolve as many potential errors as possible. There are different approaches to solving these challenges and the result is that different assemblers—when given the same sequence reads as input—will produce different assemblies.

Evaluations of genome assembler performance show that even within a phylum, there is no single optimum: the Assemblathon2 competition used 102 metrics to compare assemblies from three chordate species (Bradnam et al., 2013); within the molluscs genome size dictated the choice of preferred assembler (Sun et al., 2021); and others have demonstrated the challenges of constructing a haploid assembly from tetraploid protozoa and plants (Pollo et al., 2020; Jung et al., 2020). There studies, and many others, demonstrate the need to continually evaluate assemblers for one’s favourite species, which for us are the nematodes.

The phylum Nematoda is home to species that parasites of humans, livestock, and crops, free-living contributors to environmental health, and pivotal model organisms that help in the understanding of human biology. Accordingly, nematodes have often led the way in animal genomes: the first animal species to have its genome sequenced was *Caenorhabditis elegans* (*C. elegans* Sequencing Consortium, 1998); at least 177 genome assemblies—generated from a broad range of assemblers—are available for at least 135 species (WormBase Parasite release 18) (Howe et al., 2017). This availability of genomic information has led to comparative genomic studies on various scales, much of it focused on the evolution of gene families (Wasmuth et al., 2008; Coghlan et al., 2019; Baker et al., 2021). For this present study, in addition to understanding how different genome assemblers perform with different nematode species, we wanted to look deeper at the downstream consequences of the technical variation. It has been shown that the number of apparent lineage-specific genes is inflated if datasets are generated by different gene finding algorithms (Weisman, Murray & Eddy, 2022), motivated us to examine the effect of genome assembler heterogeneity on gene family comparisons.

We examined assemblies from three nematode species—*Caenorhabditis bovis*, *Haemonchus contortus*, and *Heligmosomoides bakeri*—which reflect different genomic features—size and heterozygosity—and sequencing platforms—Oxford Nanopore Technologies (ONT) and Pacific Biosciences (PacBio).

*C. bovis* is associated with bovine parasitic otitis disease and is the only pathogenic member of its genus (Kreis, 1964). Its genome assembly is approximately 63Mb, making it the smallest *Caenorhabditis* and one of the smallest nematode genomes so far sequenced (Stevens et al., 2020). *H. contortus* parasitizes small ruminants (sheep and goats) and the emergence of drug resistant strains motivated the study of its genome to understand mechanisms of resistance and develop new drugs and vaccines (Laing et al., 2013). The *H. contortus* genome assembly has undergone multiple rounds of improvement—a long-read assembly was scaffolded with optical mapping and manually curated—to give seven scaffolds, with a total length of ∼283Mb (Doyle et al., 2020). Finally, *H. bakeri* is an intestinal parasite of mice that serves as a model for studying host-parasite interactions and testing anthelmintic efficacy (Hu et al., 2013; Colomb et al., 2024). Note that earlier descriptions of the *H. bakeri* genome, were described as *Heligmosomoides polygyrus*, a long-standing miss-classification of the study species which has been recently corrected through genomic comparisons (Chow et al., 2019; Stevens et al., 2023).

We generated genome assemblies for each species using several popular genome assemblers and annotated these assemblies with a single, widely used pipeline. We found that most programs returned assemblies with encouraging and similar assembly metrics—size, number of scaffolds, gene completeness—but intra-species comparisons revealed structural rearrangements in the scaffolds and diverse complements of conserved orthologues. We explored the effect of assembly differences on annotation more deeply by carrying out a phylogenomic comparison of large gene families—the cytochrome P450s (CYP), glutathione S-transferases (GST), UDP glucuronosyltransferases (UGT), and nuclear hormone receptors (NHR)—and found that despite having the same sequence reads as input, the assemblies’ annotations could vary by as much as 51%.

## Methods

### Nematode species and sequence data

DNA sequence reads for *C. bovis* and *H. contortus* were retrieved from the Sequence Read Archive, and their genomes downloaded from WormBase Parasite (release 17) (Leinonen, Sugawara & Shumway, 2011; Howe et al., 2017). The sequence reads and assembly for *H. bakeri* are new for this study. *C. bovis* DNA (BioProject: PRJEB34497) was sequenced using the Oxford Nanopore MinION platform (Stevens et al., 2020). *H. contortus* DNA (BioProject PRJEB506) was sequenced using the PacBio RS and Sequel platforms (Doyle et al., 2020). We generated *H. bakeri* DNA as part of a *de novo* sequencing project. Briefly: we extracted adult *H. bakeri* worms from experimentally infected mice; separated females and males; pooled 20 worms of each sex. PacBio Hi-Fi sequencing was done at the University of Delaware DNA Sequencing & Genotyping Center. For the assemblies in this study, we combined sequence reads from female and male worms. The sequence reads are available from the NCBI SRA: SRX23938287 and SRX23938287.

### Genome assembly and quality control

#### Generation of preliminary assemblies

For *C. bovis* and *H. contortus* we used six long-read assemblers: Redbean v2.5 (Ruan & Li, 2020), Canu v2.1.1 (Koren et al., 2017), Flye v2.8.2 (Lin et al., 2016; Kolmogorov et al., 2019), SMARTdenovo v1.0.0 (Liu et al., 2021), Falcon-kit v1.8.1 and Falcon-unzip v1.3.7 (Chin et al., 2016). For *H. bakeri*, we used three HiFi-compatible assemblers: Flye v2.8.2 (Lin et al., 2016; Kolmogorov et al., 2019), Hifiasm v0.16.1-r375 (Cheng et al., 2021) and HiCanu v2.1.1 (Nurk et al., 2020). All the commands and parameters can be found in File S1.

#### Decontamination and polishing of preliminary assemblies

For the *C. bovis* and *H. contortus*, we used Blobtools v1.1.1 (Laetsch & Blaxter, 2017). Long reads were aligned to assembly contigs using minimap2 v2.17-r941 (Li, 2018) and the taxonomic classification determined by searching contigs against the Uniprot Proteome database using DIAMOND v2.0.6.144 (Buchfink, Xie & Huson, 2015; Pundir, Martin & O’Donovan, 2017). For polishing—identification and correction of small errors—the *C. bovis* and *H. contortus* assemblies underwent multiple rounds of polishing using tools appropriate for the sequencing platforms used. Briefly:

1. Four rounds with Racon with long reads for both *C. bovis* and *H. contortus* assemblies (Vaser et al., 2017),
2. One round with Medaka v1.3.2 (Oxford Nanopore Technologies., 2018) with long reads for *C. bovis* assemblies (https://github.com/nanoporetech/medaka),
3. One round with Arrow v2.3.3 with long reads for *H. contortus* assemblies (installed via the GenomicConsensus package v2.3.3, http://github.com/PacificBiosciences/pbbioconda),
4. Two rounds of Racon and Pilon v1.14 Assemblies with Illumina reads for both *C. bovis* and *H. contortus* assemblies (Walker et al., 2014; Vaser et al., 2017).

All the commands and parameters can be found in File S1.

#### Removal of residual haplotype duplications

This was done for all *H. contortus* and *H. bakeri* assemblies generated as part of this study. We used the Purge_dups v1.2.5 tool (Guan et al., 2020). As recommended in the Purge_dups documentation, we initially applied the default cut-off threshold, followed by two manually set cut-offs derived from the default cutoffs (Table S10). All the commands and parameters can be found in File S1.

#### Assembly evaluation

For all three species, all the generated assemblies were evaluated for contiguity and completeness using various metrics including: number of fragments, assembly size, N50, length of the longest contig, and BUSCO v4.1.4 (Simão et al., 2015). All the commands and parameters can be found in File S1. Briefly, we used BUSCO v4.1.4 to evaluate completeness of single copy orthologues (Manni et al., 2021) and Inspector v1.0.2 to identify both large- and small-scale errors (Chen et al., 2021).

### Genome synteny

We used Nucmer (from MUMmer v3.23) to create assembly-to-assembly alignments (Marçais et al., 2018). We used the following assemblies as the reference for Nucmer: Redbean v2.5 of *C. bovis*; Doyle for *H. contortus*; Hifiasm for *H. contortus*. For *C. bovis*, we used the dnadiff script (part of Nucmer) to report one-to-one alignment coordinates and viewed the alignments with Cirocs v0.69-8 (Krzywinski et al., 2009). For all species, we used NucDiff v2.0.3 to measure large- and small scale differences between each assembly and the reference (Khelik et al., 2017). All the commands and parameters can be found in File S1.

### Visualising BUSCO content analysis

We generated Sankey diagrams using the SankeyMATIC builder (available at http://sankeymatic.com/build/) to visualize the transitions of BUSCO genes, i.e., those genes that changed categories between assemblies. Set-based BUSCO genes data was displayed using UpSet plots (Lex et al., 2014). For *H. contortus*, we created a heatmap of missing BUSCO genes with the ggplot2 package in R v3.4.2 (Wickham, 2016; R Core Team, 2022). Most figures were edited with Adobe Illustrator (available at https://www.adobe.com). Whenever possible, a colour-blind accessible palette was chosen using (https://davidmathlogic.com/colorblind/).

### Functional annotation of BUSCO genes

We used BlastKOALA webserver (Kanehisa, Sato & Morishima, 2016).

### Genome annotation

#### Soft masking assemblies

We used RepeatModeler v2.0.3 and RepeatMasker v4.1.2-p1 (Tarailo-Graovac & Chen, 2009; Flynn et al., 2020). For RepeatModeler, we used NCBI/RMBLAST v2.11.0+ to create repeat library from the Dfam database of transposable elements and DNA repeats (Storer et al., 2021). Next, we used the RepeatMasker util (queryRepeatDatabase.pl script) to create a custom library specific to nematode species. Both repeat libraries were combined as input for the RepeatMasker program.

#### Gene prediction

Protein coding genes (gene models) in the generated assemblies were predicted using Braker 3 (Gabriel et al., 2023) installed using the Singularity container (available at https://hub.docker.com/r/teambraker/braker3). For *H. contortus* and *H. bakeri*, we used short-read RNA-seq data from the Sequence Read Archive as external evidence for the gene prediction (BioProjects: PRJEB506 (Laing et al., 2013) and PRJNA750155 (Pollo et al., 2023) respectively). RNA-seq data was aligned using the STAR v2.7.10a aligner (Dobin et al., 2013) and the generated alignments, in bam-format, used to produce a training set for AUGUSTUS v3.5.0, using GeneMark-ET v4.71 (Hoff et al., 2016). We filtered redundant training gene structures with DIAMOND v0.9.24 (Buchfink, Xie & Huson, 2015). For *C. bovis*—for which no RNA-Seq is available—we used the longest isoforms from *C. elegans* gene models (BioProject: PRJNA13758) retrieved from WormBase Parasite (release 17) (Howe et al., 2017), and the default protein sequences retrieved from UniProtKB/Swiss-Prot database (Boutet et al., 2007) as the protein evidence for gene prediction. Braker 3 used the ProtHint pipeline to produce hints using GeneMark-EP v4.71, DIAMOND v0.9.24 and Spaln v2.3.3d tools (Lomsadze et al., 2005; Iwata & Gotoh, 2012; Gotoh, Morita & Nelson, 2014; Buchfink, Xie & Huson, 2015; Brůna, Lomsadze & Borodovsky, 2020). All the commands and parameters can be found in File S1.

#### Identification of gene family members

Members of the Cytochrome P450 (CYP), Glutathione S-transferase (GST), UDP-glucuronosyltransferase (UGT) and nuclear hormone receptors (NHR) gene families were identified using BLASTP v2.9.0+ against the full *C. elegans* protein set (Camacho et al., 2009). For each species, we clustered the proteins for each family using OrthoFinder2 (‘-S blast’, all other parameters were default) and included the *C. elegans* proteins to serve as assumed outgroups (Emms & Kelly, 2019).

For the TGM protein family, the protein sequences were taken from the publication (Smyth et al., 2018). The latest ARI and BARI protein sequences from provided by Dr. Henry McSorley (University of Dundee) (Osbourn et al., 2017). These protein sequences were mapped to the genome assemblies using the splice-aware aligners exonerate and miniprot (Slater & Birney, 2005; Li, 2023). For both, we a scoring threshold which only accepted an alignment which scored within a specified percentage of the top score for a given query. For miniprot we used ‘--outs 90’, and for exonerate we used ‘--percent 90’ and ‘--percent 70’. We found that the results for ‘exonerate --percent 70’ were broadly similar to ‘miniprot --outs 90’ (Table S69).

## Results

We generated fifteen new assemblies—six for *C. bovis*, six for *H. contortus*, and three for *H. bakeri*—which were used, along with the published genomes for each species, for intraspecies comparisons. All the assemblies are available in File S2. For readability, we will refer to the published genomes by their papers’ first author— Stevens for *C. bovis*, Doyle for *H. contortus*, and Chow for *H. bakeri*—and by the name of the genome assembly program for the others (Chow et al., 2019; Stevens et al., 2020; Doyle et al., 2020). Each new assembly was subject to decontamination and polishing protocols with an additional step of purging duplicates in the highly heterozygous *H. contortus* and *H. bakeri* assemblies. Decontamination—the removal of contigs from bacteria and other heterogenous sources—removed approximately 20% of the nucleotides of any given intermediate assembly and, in *C. bovis*, up to 90% of the scaffolds (Tables S7-S9). Polishing—using the sequence reads to identify and fix any small errors—led to notable improvements in the ONT *C. bovis* assemblies; for example, the Redbean2.5 assembly was improved from 79% to 96% of the BUSCO genes were found as complete (Table S7). However, polishing had a minor impact on the PacBio-based assemblies, *H. contortus* (CSII) and *H. bakeri* (HiFi), presumably because the genome assemblers used had their own in-built polishing steps. Purging duplicates— phasing out allelic repeats in highly heterozygous genomes based on read depth—led to notable reductions in the genome sizes and duplication of BUSCO genes; for example, the BUSCO genes’ duplication in the *H. contortus*’ Canu assembly was reduced by 48% (Table S8).

Below, we compare the assemblies of each species using general assembly statistics, synteny, and BUSCO gene completeness. We then use four gene families to demonstrate how differences in assembly impact downstream analysis.

### *C. bovis* assemblies

The Stevens *C. bovis* assembly was generated using ONT reads with the Redbean assembler v2.3 and polished with Racon and Medaka. We generated assemblies using an updated Redbean (v2.5, hereafter Redbean2.5), Flye, Canu, SMARTdenovo, Falcon, and Falcon-Unzip (Chin et al., 2016; Lin et al., 2016; Koren et al., 2017; Kolmogorov et al., 2019; Ruan & Li, 2020; Liu et al., 2021). Following decontamination and polishing (Table S7), the overall standard assembly metrics were relatively similar (Table 1). Compared to the Stevens assembly, the Redbean2.5 assembly was near identical in size but on fewer scaffolds (21 vs 35) and the N50 was 400 kb larger. The Canu assembly size was 10% larger and on almost four times as many scaffolds. The other four assemblies were comparable to the Stevens assembly in terms of assembly size and the number of scaffolds, but the N50s were between 25% and 50% smaller.

**Table 1:** Assembly statistics for *Caenorhabditis bovis.* Complete ‘C’ gene the sum of single copy ‘S’ and duplicate ‘D’ (all % from 3131 genes).

| Assembly Stats | Stevens | Redbean v2.5 | Flye | Canu | SMARTdenovo | Falcon | Falcon-unzip |
| --- | --- | --- | --- | --- | --- | --- | --- |
| # Scaffolds | 35 | 21 | 34 | 134 | 32 | 48 | 41 |
| Assembly size (Mb) | 62.73 | 62.72 | 62.93 | 69.16 | 63.39 | 64.93 | 64.40 |
| N50 (Mb) | 7.60 | 8.03 | 5.86 | 4.01 | 3.56 | 4.41 | 4.41 |
| Longest contig (Mb) | 10.86 | 10.08 | 10.12 | 12.16 | 6.29 | 9.90 | 9.79 |
| BUSCO %<br>n=3131 | C:96.2<br>S:95.6 D:0.6 | C:96.0<br>S:95.3 D:0.7 | C:96.4<br>S:95.5 D:0.7 | C:96.3<br>S:93.2 D:3.1 | C:96.1<br>S:93.2 D:0.9 | C:96.1<br>S:95.0 D:1.1 | C:96.2<br>S:95.1 D:1.1 |
| Inspector QV | 25.1 | 25.1 | 25.0 | 25.3 | 25.1 | 25.0 | 25.2 |
| Structural errors | 12 | 8 | 9 | 10 | 16 | 21 | 9 |

Through BUSCO analysis, we found a similar percentage of genes were complete across the seven assemblies (Table 1 and Tables S11-S17). A notable difference was the duplication rate in the Canu assembly, which was five times that of the Stevens assembly: 18 genes vs 97 genes.

Of the 3131 BUSCO orthologues, 3040 genes (97%) were complete (single copy or duplicated) in at least one assembly and 94.9% (2972) were complete in all seven assemblies (Figure S1). Of these complete genes, 15 were duplicated in all assemblies. Ninety-one genes were missing or fragmented in all seven assemblies, which suggests they are undergoing pseudogenization or have been lost as a consequence of a genome size reduction in *C. bovis* (Stevens et al., 2020). Ninety-one (3%) were absent (fragmented or missing) in all seven assemblies. Six genes were complete in only one assembly. In total, the BUSCO classification differed for 178 genes, *e.g.* from Duplicated to Fragmented (Figure 1A). Expectedly, there was strong agreement between the Stevens (Redbean2.3) and the new Redbean2.5 assemblies; nine genes complete and single in the Stevens assembly were fragmented or missing in the Redbean2.5 assembly with six genes in the reverse situation (Figure 1A). Disagreements on BUSCO classification was more striking between the other assemblies. For example, SMARTdenovo and Falcon had near identical BUSCO scores, but 36 single copy genes in the SMARTdenovo assembly were classified as duplicated (16), fragmented (13) or missing (7) in the Falcon assembly. In a direct comparison of Redbean2.5 and Canu assemblies, 18 genes considered fragmented or missing in Redbean2.5 but were found complete and single copy in Canu (Figure 1B).

**Figure 1:**
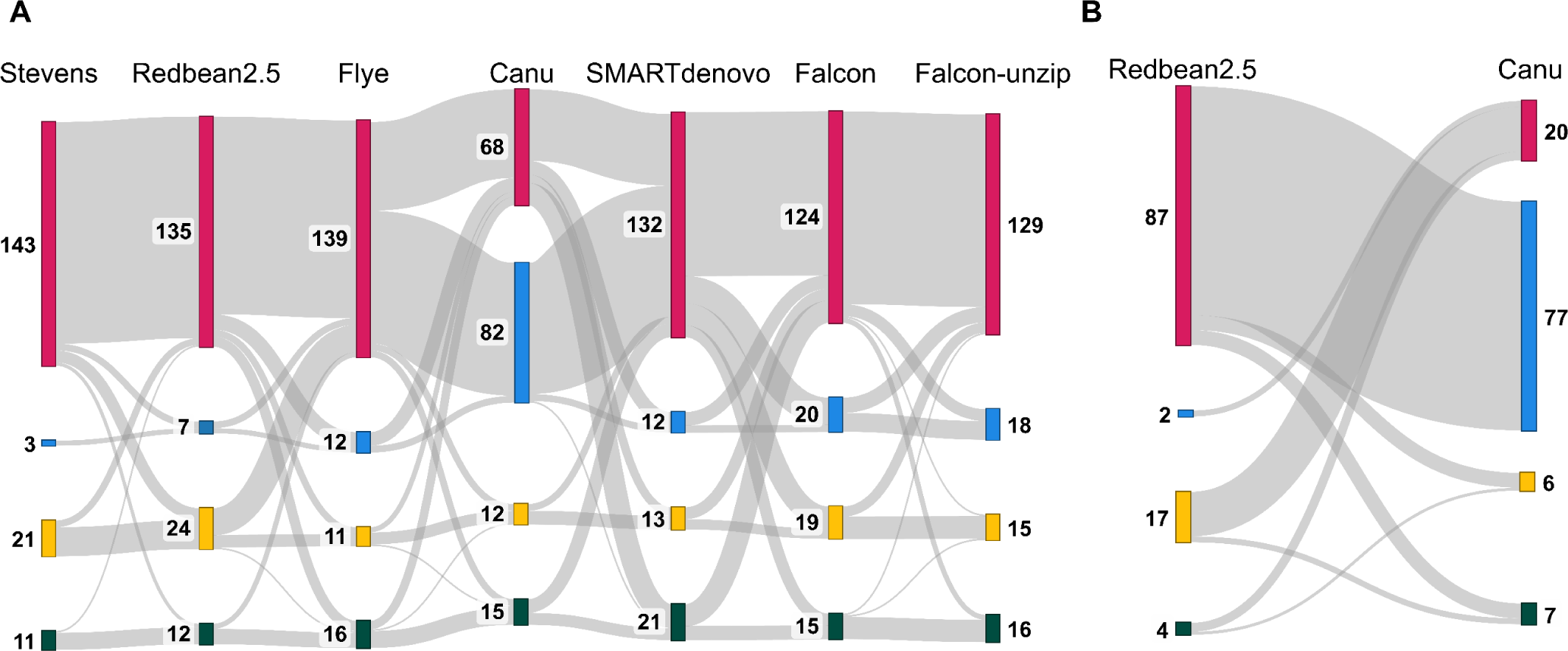
Transitions of BUSCO genes across multiple *C. bovis* assemblies. The Sankey diagrams show how BUSCO genes switch categories: red – complete and single copy genes; blue – complete and duplicated; yellow – fragmented; green – missing. (A) The 178 genes that changed BUSCO categories in at least one assembly. The order of the assemblies is arbitrary and follows Table 1. The data for this panel is in Table S1 and Table S2. (B) Comparison of the 110 genes which differed between Redbean2.5 and Canu assemblies. The data for this panel is in Table S3.

We used the original ONT reads to evaluate the reads alignment to each of the assemblies, identifying large structural and small errors (Chen et al., 2021). All the assemblies had near identical QV scores, the Redbean2.5 assembly had the fewest large structural errors, and the Flye assembly had the fewest small-scale errors (Table 1 and Table S18).

Next, we investigated the structural variation—insertions, deletions, and translocations—between assemblies. We needed a reference assembly, and considered Redbean2.5 assembly to have the best accuracy metrics, albeit slightly. As expected, the Stevens and Flye assemblies—both use the DBG algorithm—had the fewest number of differences (Table 2 and Table S19). These were followed by the OLC-based SMARTdenovo assembly, the related HGAP Falcon and Falcon-Unzip assemblies, with the OLC-based Canu assembly having the most differences to the Redbean2.5 assembly. For all assemblies, at least 97% of the variants were insertions or deletions of typically short regions (1-to-10bp). However, we did find longer duplications and tandem duplications which could impact accurate curation of genes. Using Braker3 to annotate the assemblies with gene models, we found five genes from the Stevens assembly—four genes in tandem duplications and one gene on a duplication—and 19 genes from the Redbean2.5—17 genes in tandem duplications and two on duplications—in these regions (Table S20).

**Table 2:** NucDiff results for *C. bovis* assemblies. Each assembly was aligned to the reference assembly, Redbean2.5.

|  | Stevens | Flye | Canu | SMARTdenovo | Falcon | Falcon-unzip |
| --- | --- | --- | --- | --- | --- | --- |
| Total number | 8,215 | 8,527 | 62,971 | 12,895 | 19,319 | 16,540 |
| Insertions | 2,421 | 3,350 | 25,809 | 4,913 | 8,002 | 6,718 |
| Deletions | 5,553 | 4,916 | 36,399 | 7,684 | 10,929 | 9,363 |
| Translocation | 194 | 207 | 478 | 218 | 290 | 336 |
| Relocations | 29 | 18 | 142 | 38 | 39 | 57 |
| Reshufflings | 8 | 11 | 32 | 15 | 21 | 18 |
| Inversions | 10 | 25 | 109 | 27 | 38 | 48 |
| Unaligned seq | 0 | 0 | 2 | 0 | 0 | 0 |

Larger structural variants—translocations, relocations, reshufflings, and inversions—were present even between Redbean assemblies; we found that the RedBean2.5 Scaffold 3 was split across the Stevens Scaffolds 6, 7, and 11 (Figure 2). However, for five of the assemblies, all scaffolds could be aligned to the reference. The exception was the Canu assembly; two scaffolds did not align to the Redbean2.5 assembly. Through a BLASTX search against NCBI NR database (Camacho et al., 2009), we found that these were most likely of bacterial origin and were not removed by the decontamination protocols.

**Figure 2:**
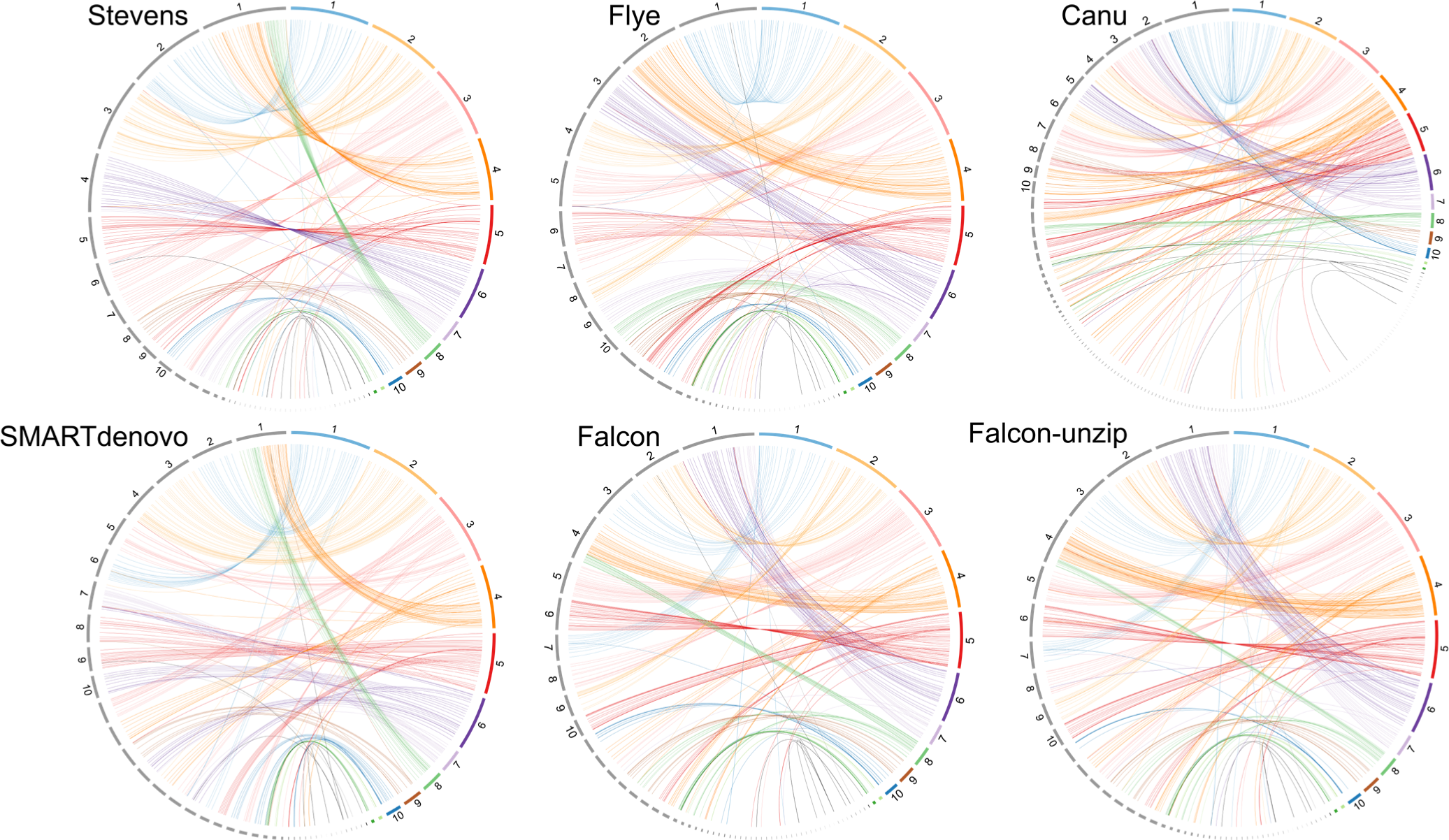
Synteny across six genome assemblies for the *C. bovis*. In each Circos plot, the reference assembly (Redbean2.5) is represented by the coloured scaffolds and the labeled assembly is represented by the gray scaffolds. The lines linking syntenic regions are coloured according to the reference scaffold. The scaffolds are ordered by length (longest to shortest) and for clarity we only label the first ten scaffolds.

### *H. contortus* assemblies

The published *H. contortus* assembly is the result of an ongoing multinational consortium and involved deep sequencing using multiple platforms and labour-intensive manual identification and correction of assembly errors (Doyle et al., 2020). We generated six assemblies using only the PacBio reads that were used in Doyle assembly. These six assemblies, once decontaminated and polished, varied considerably in their length, fragmentation, and completeness (Table 3). There was a 23% difference between the longest (Flye 331 Mb) and shortest (Falcon-unzip 262 Mb) assemblies. The SMARTdenovo assembly was the most contiguous, with the fewest scaffolds and largest N50. The Redbean2.5 and Falcon assemblies were the closest to the Doyle assembly in terms of assembly length.

**Table 3:** Assembly statistics for *H. contortus*. Complete ‘C’ gene the sum of single copy ‘S’ and duplicate ‘D’ (all % from 3131 genes).

|  | Doyle | Redbean v2.5 | Flye | Canu | SMARTdenovo | Falcon | Falcon-unzip |
| --- | --- | --- | --- | --- | --- | --- | --- |
| # Scaffolds | 7 | 1844 | 3249 | 2364 | 1473 | 2402 | 1975 |
| Assembly size (Mb) | 283.44 | 296.37 | 331.35 | 341.72 | 314.72 | 267.14 | 262.37 |
| N50 (Mb) | 47.40 | 0.31 | 0.24 | 0.34 | 0.45 | 0.23 | 0.25 |
| Longest contig (Mb) | 51.80 | 3.45 | 4.85 | 2.76 | 4.47 | 3.22 | 3.24 |
| BUSCO %<br>n=3131 | C:94.8<br>S:93.7 D:1.1 | C:93.9<br>S:86.7 D:7.2 | C:94.1<br>S:85.0 D:9.1 | C:95.2<br>S:86.6 D:8.6 | C:94.7<br>S:86.5 D:8.2 | C:88.2<br>S:84.2 D:4.0 | C:87.7<br>S:84.2 D:3.5 |
| **Inspector Reference-free (mapping reads-to-assembly):** |  |  |  |  |  |  |  |
| Inspector QV | 22.0 | 42.9 | 39.6 | 44.6 | 39.1 | 31.8 | 42.8 |
| Structural errors | 170 | 53 | 146 | 67 | 180 | 826 | 62 |
| **Inspector results Reference-guided (mapping assemblies-to-Doyle):** |  |  |  |  |  |  |  |
| Coverage of Doyle (%) | 98 | 84 | 80 | 84 | 84 | 79 | 79 |
| Reference bases with depth<br>0; 1; 2+ (%) | 2; 98; 0 | 16; 82; 2 | 20; 76; 4 | 16; 78; 6 | 16; 81; 3 | 21; 77; 2 | 21; 77; 2 |
| Number of collapses | 0 | 1197 | 1633 | 1512 | 1234 | 6370 | 1019 |
| Number of expansions | 0 | 1004 | 1136 | 1127 | 1113 | 845 | 888 |
| Number of inversions | 0 | 1 | 0 | 0 | 0 | 2 | 0 |

For the Inspector reports, we used both the reads and the Doyle assembly for reference-free and reference-guided metrics, respectively (Table 3 and Table S21). We were initially surprised to see that the Doyle assembly had the lowest QV score. Here, it is important to note that this score is reference-free, and a function of read alignments to the assembly. The Doyle assembly has undergone extensive curation, in which data, including optical mapping, was used to identify mistakes in the preliminary long-read driven assembly. So, a higher level of disagreement with the raw sequence reads might be expected. For example, the Doyle assembly had the highest number of haplotype switches likely a consequence of the manual improvement that sought to maximize scaffold continuity (Table S21). For the six assemblies that we generated, the Inspector metrics were similar. The coverage of the assembly-to-Doyle alignments were relatively low (79%-to-84%). The effect of the alignment algorithm is demonstrated with the 98% coverage Doyle-to-Doyle assembly. Therefore, the regions not covered predominantly are due to differences between each of the six assemblies and the Doyle reference. The duplication rate—where the assembly-to-Doyle coverage was two or more—was lowest for the Redbean, Falcon and Falcon-unzip assemblies and highest in the Canu assembly.

In considering the BUSCO results, the six new assemblies varied more than those of the other two species, with between 87.7% to 95.2% of genes found complete. The duplication rate was notably higher in the six assemblies compared to the Doyle reference. Of the 3131 pan-nematode orthologous genes, 3020 genes (96.5%) were complete in at least one assembly, with 2525 genes (80.6%) complete in all seven assemblies and 94 genes (3.0%) missing or fragmented in all seven (Figures S2-S4). It is striking that fewer genes were missing in the Canu and SMARTdenovo assemblies compared to the Doyle assembly. We looked more broadly and found that of the 141 genes missing in the Doyle assembly, 56 (39.7%) were complete at least one of the new assemblies, 27 in all six assemblies (Figure 3). It is likely that these represent errors in the current reference assembly.

**Figure 3:**
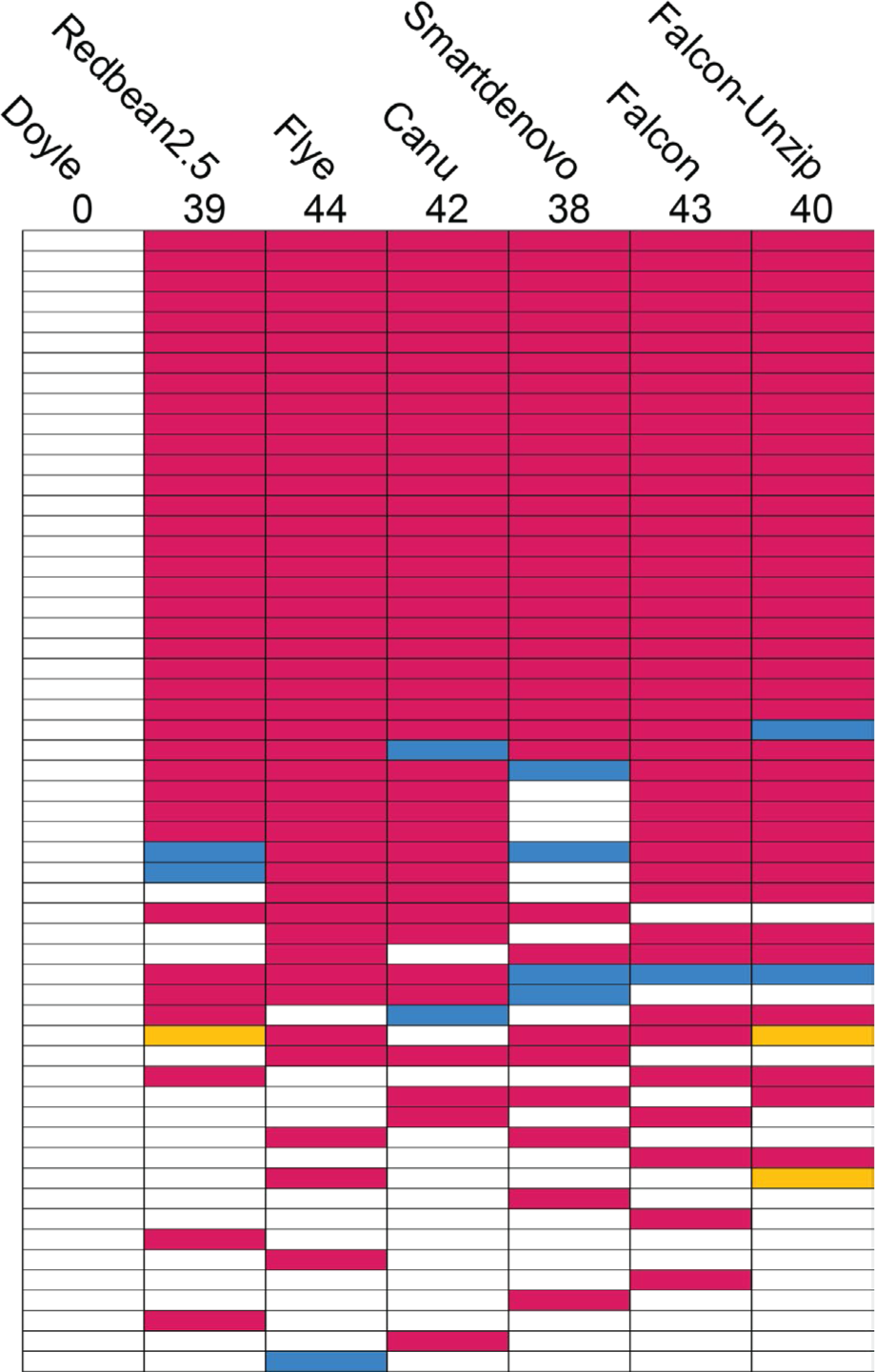
BUSCO genes missing from the *H. contortus* Doyle assembly but complete in at least one other assembly. Red – complete and single copy genes; blue – complete and duplicated; yellow – fragmented; white – missing. The data for this figure is in Table S6.

We used BlastKOALA to annotate these 141 genes and found at least three noteworthy examples (Table S22) (Kanehisa, Sato & Morishima, 2016). One example is *let-756* which is involved in the fibroblast growth factor receptor signaling pathway essential in nematode larval development. The knock-down of *let-756* in *C. elegans* by RNA interference (RNAi) induces larval arrest in worms (Roubin et al., 1999). A second example is *nck-1*, which enables kinase activity required for neuronal guidance. The knock-down of *nck-1* in *C. elegans* negatively disrupts in axon guidance, neuronal cell positioning, and abnormalities in the excretory canal cell, gonad, and male mating (Mohamed & Chin-Sang, 2011). A third example is *gcn-2*, a kinase involved in the nutrient-sensing and, while knock out is not fatal in *C. elegans*, it does severally impact *C. elegans* survival during nutrient stress, specifically amino acid limitation (Rousakis et al., 2013).

Of the 35 BUSCO genes duplicated in the Doyle reference assembly, 29 were also duplicated in most of the new assemblies and three were single copy in all six new assemblies (Figure S2). There were large exchanges between single copy and duplicated categories, suggesting that assembly algorithms successfully collapsed divergent haplotypes of different genes at different rates (Figure S5 and Table S4). The presence or absence of BUSCOs between the new assemblies was striking. For example, 44 genes considered missing or fragmented in the Doyle reference assembly (‘red’) were classified as complete and single copies in the Redbean2.5 assembly (Figure S5). Another example is, 62 genes considered missing or fragmented in the Redbean2.5 assembly single copy in Flye (Figure S5), with 66 genes reclassified in the other direction (Figure S6 and Table S5). Both assemblers use the DBG algorithm. The full list of BUSCO genes for each assembly can be found in Tables S23-S29).

### *H. bakeri* assemblies

At the start of our study, there were two published *H. bakeri* assemblies; one generated from short-reads only and one that used a low-coverage of PacBio reads to scaffold together a short-read assembly (Coghlan et al., 2019; Chow et al., 2019). We selected the Chow assembly as it was more contiguous and had a higher BUSCO score. As we finished our study, a chromosome-scale assembly for *H. bakeri* was published (Stevens et al., 2023). While more contiguous than the Chow assembly, we note that the Chow assembly had a higher BUSCO score (Table 4).

**Table 4:**
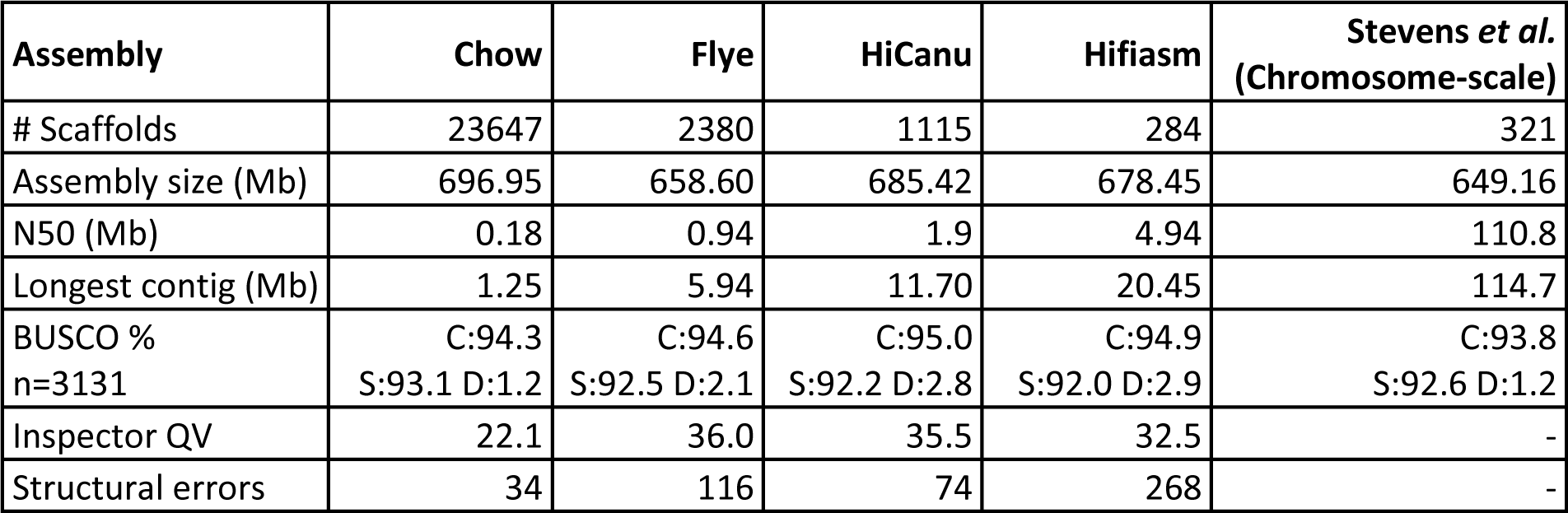
Assembly statistics for *H. bakeri*. Complete ‘C’ gene the sum of single copy ‘S’ and duplicate ‘D’ (all % from 3131 genes). The chromosome-level Stevens *et al*. assembly was only published as we completed our study and was not included in our comparisons.

As part of our own *H. bakeri* genome project, we generated HiFi PacBio reads and assembled them with three different software: Flye, HiCanu, and Hifiasm. The assembly lengths differed less than 6%, but there was considerable variation of N50 and longest contig (Table 4).

The three newly generated assemblies exhibited comparable performance in the reference-free Inspector analysis. However, the Hifiasm assembly, despite its higher contiguity, exhibited more structural errors (Tables 4 and S30).

Similar to *C. bovis* and *H. contortus*, the *H. bakeri* assemblies, despite extremely similar BUSCO scores, contained different cohorts of BUSCO genes (Figure 4 and Table S31). A total of 120 genes were complete in three assemblies and absent (fragmented or missing) in one (Figure 5A. We note that while 58 of these in-three-out-one genes were absent in the Chow assembly, 62 genes were fragmented or missing in one of the HiFi-based assemblies. We explored the functions of these 62 genes to determine their potential impact on hypotheses regarding *H. bakeri* biology (Tables S32-S38).

**Figure 4:**
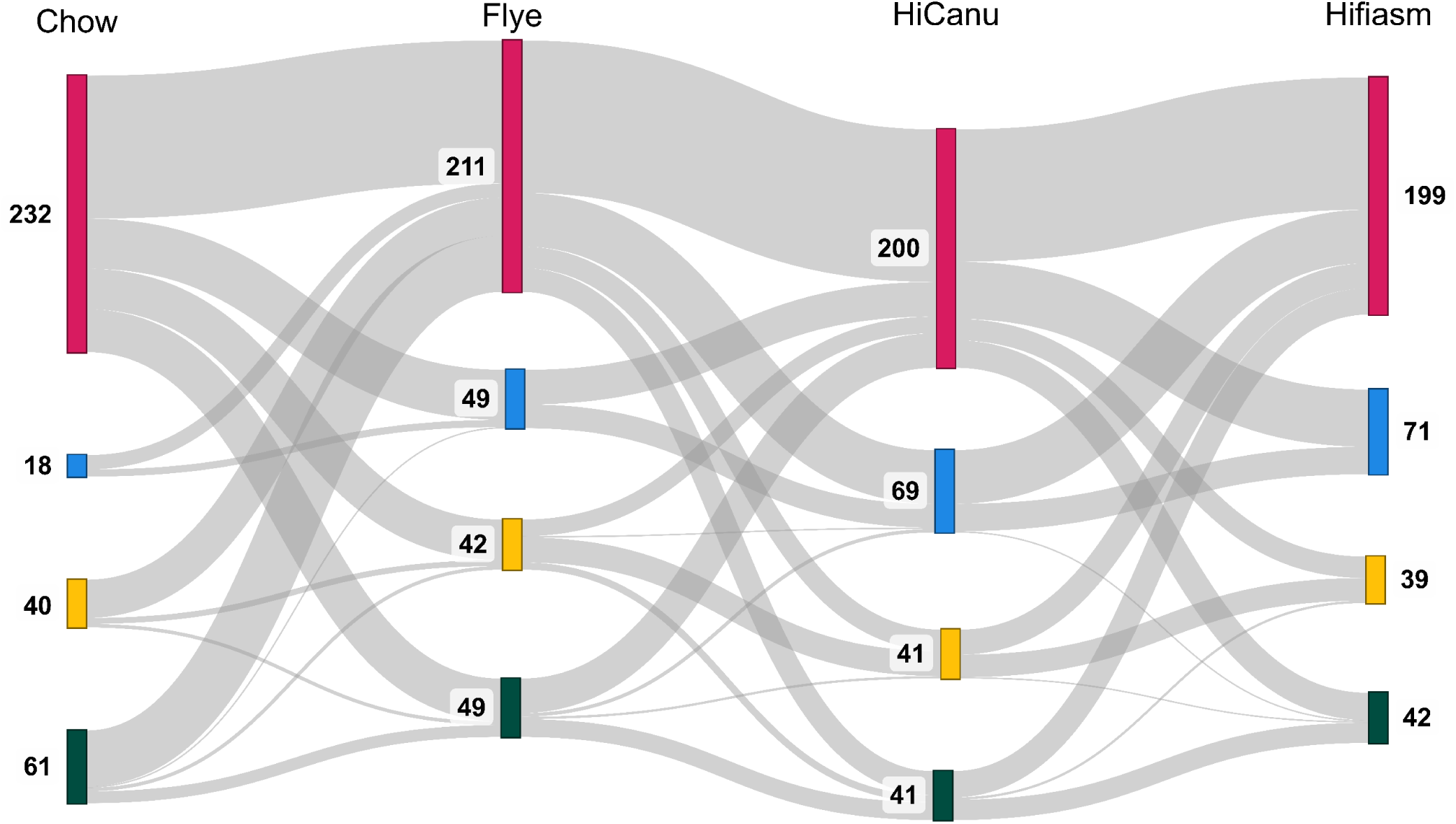
Transitions of BUSCO genes across multiple *H. bakeri* assemblies. The Sankey diagram shows how 351 genes changed categories in at least one *H. bakeri* assembly: red – complete & single copy genes; blue – complete & duplicated; yellow – fragmented; green – missing. The order of the assemblies is arbitrary and follows Table 4. The data for this figure is in Table S31.

**Figure 5:**
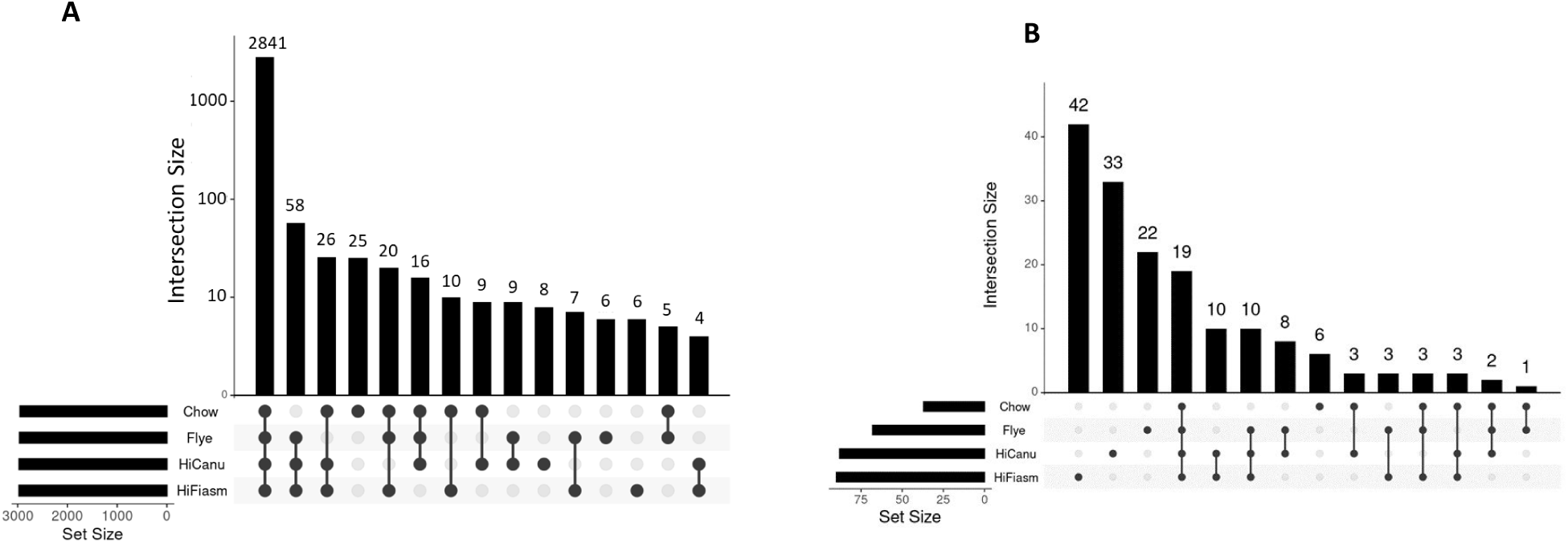
UpSet plots of (A) Complete BUSCOs present in *H. bakeri* assemblies, and (B) Duplicated BUSCOs in the assemblies. The figures were generated with https://asntech.shinyapps.io/intervene/.

We found several genes whose absence, if real, could significantly affect a *H. bakeri* worm. One example is *mogs-1*, absent only in the Flye assembly, which encodes a mannosyl-oligosaccharide glucosidase which is involved in N-glycan biosynthesis and plays a pivotal role in digestion of bacteria (Geng et al., 2022). A second example is *gpb-1,* also absent in the Flye assembly, which is a G protein beta subunit and whose RNAi knock-down is embryonic lethal (Kamath et al., 2000). A third example, absent only in the HiCanu assembly, is *fmo-4*, which encodes for flavin-containing monooxygenase crucial for osmoregulation and RNAi knock-down is also embryoic lethal (Rual et al., 2004; Hirani et al., 2016).

While, marginally more BUSCO genes were found in the HiFi assemblies compared to the Chow assembly, it came at the cost of a higher duplication. We found that 165 BUSCO genes were duplicated in at least one assembly; the majority of these were assembly specific with 19 found duplicated in all four assemblies and a further 10 found in the HiFi-based assemblies (Figure 5B). This tempts speculation that the ∼29 genes are indeed duplicated.

### Impact of assembly variation on gene family analysis

We generated protein-coding gene models for all assemblies using Braker3 (Table 5 and File S2) (Gabriel et al., 2023). For consistency, we used Braker3-generated gene models for the published genomes (see Discussion).

**Table 5:**
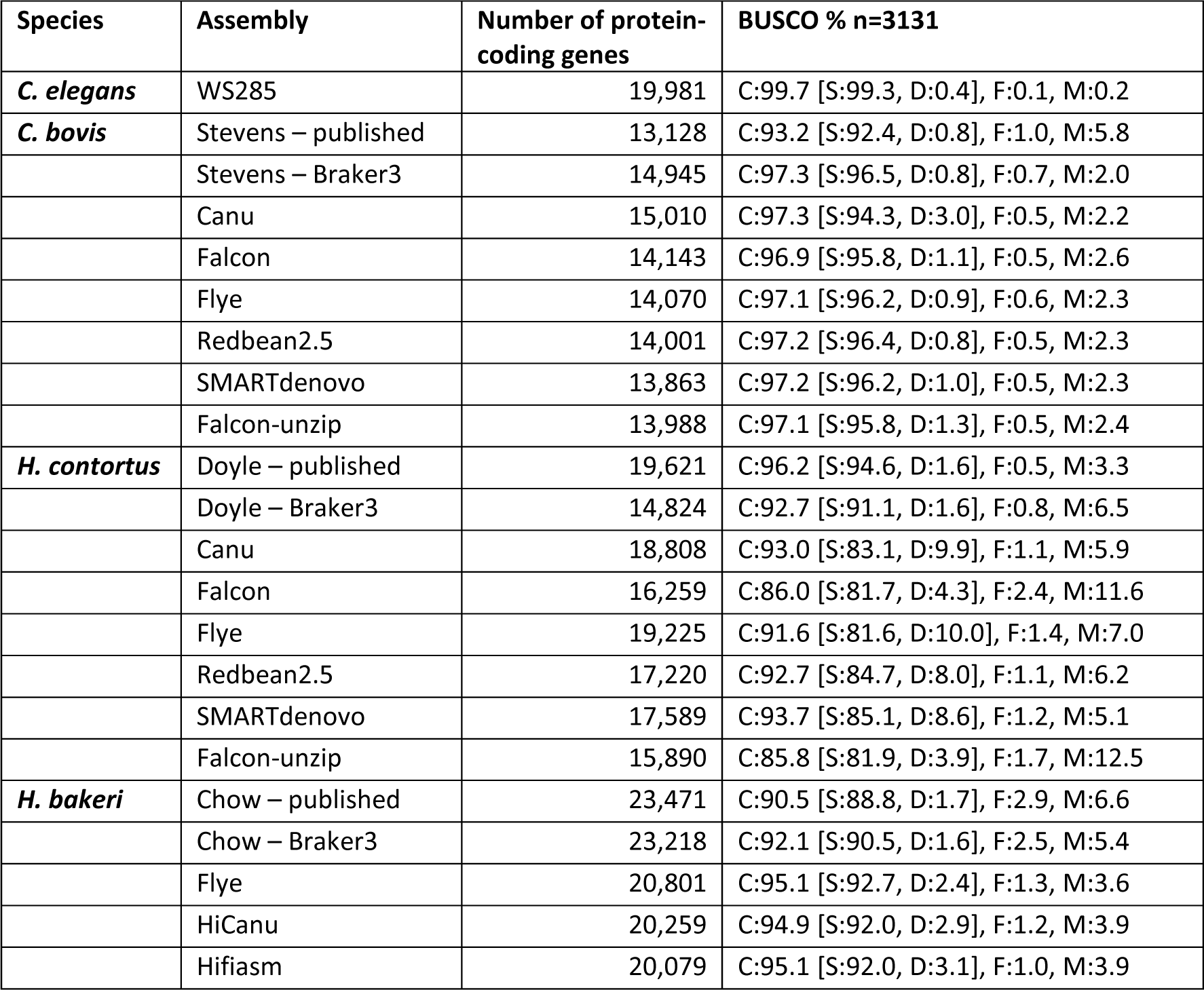
Protein-coding genes predicted for the different assemblies. For the published genomes, we report the both the published protein-coding gene models and our reannotation. For all other assemblies, the protein-coding gene models were generated with Braker3. BUSCO scores (C=complete, S=single copy, D=duplicated, F=fragmented, M=missing) are percentages where *n*=3131.

For all protein sets, we identified members of four gene families—cytochrome P450s (CYP), glutathione S-transferase (GST), UDP glucuronosyltransferases (UGT), and nuclear hormone receptors (NHR)—which have been implicated in anthelmintic studies and are among the largest in *C. elegans* (Tables S39-S43). For each gene family, within each species, we clustered the proteins with those from C. elegans (Emms & Kelly, 2019). Here, our assumption is that if the differences in genome assemblies did not affect the genome annotation, we would expect that every cluster would have an identical number of proteins from each assembly and that these would form perfect one-to-one orthologous groups to the exclusion of *C. elegans*. Any deviation from this would demonstrate that differences in the assemblies led to different gene model predictions. We considered cluster membership across all assemblies (‘all’) and just for those assemblies generated in this study (‘new’) (Table 6).

**Table 6:**
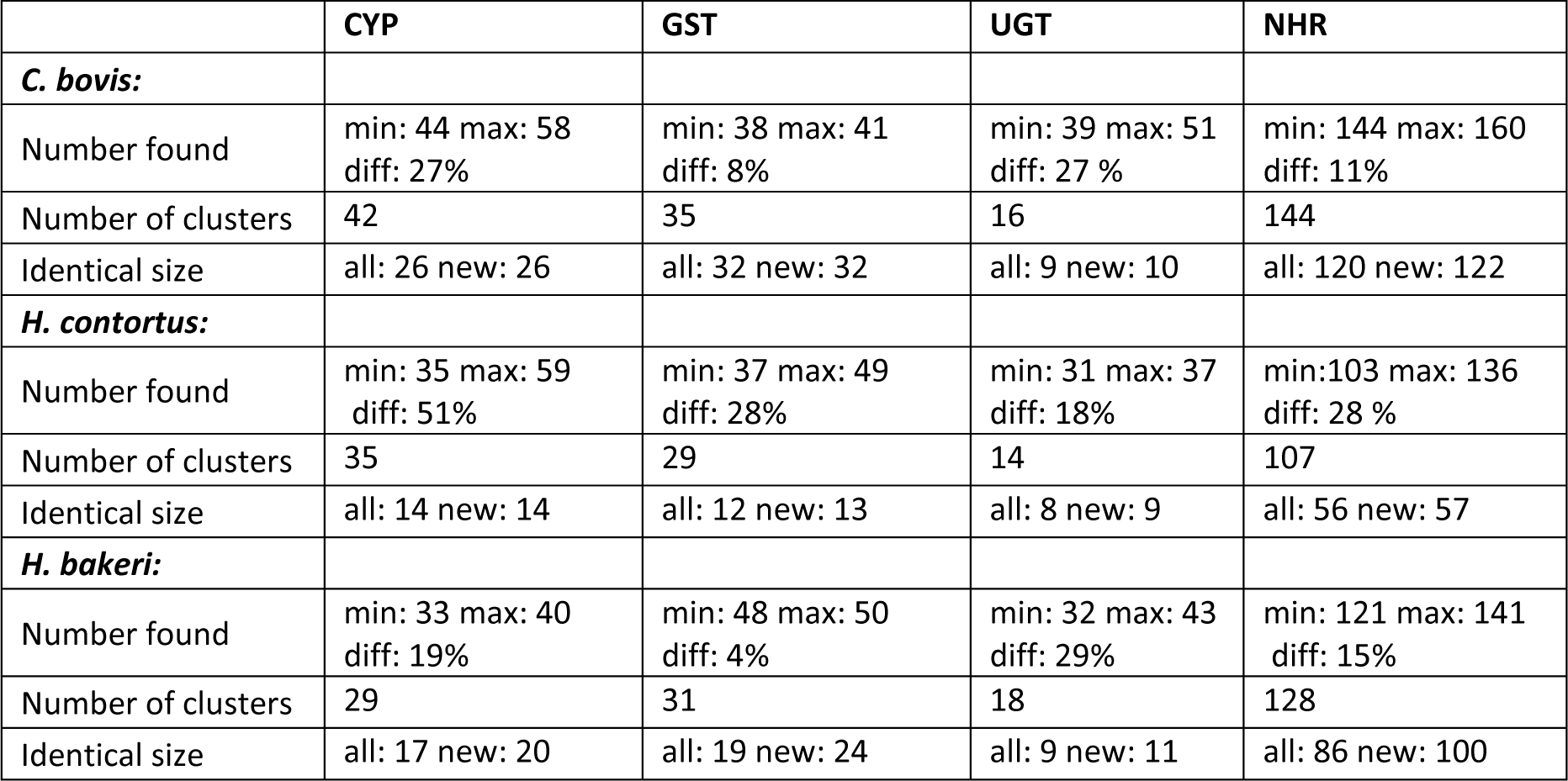
Showing percentage differences between the minimum and maximum number of the predicted members of the CYP, GST, UGT and NHR gene families across assemblies. We considered cluster membership across all assemblies (‘all’) and also restricted to those assemblies generated in this study (‘new’).

Overall, we found considerable variation within gene family membership (Table 6 and Tables S45-S68). We looked at the correlation between the size of the gene families and the size of the assembly or the BUSCO duplication rate. The results were inconsistent. We saw a strong and positive correlated (ρ>0.7) for: GSTs, UGTs, and NHRs in *C. bovis* against both assembly size and duplication rate; and NHRs in *H. contortus* against both assembly size and duplication rate. However, we saw a strong and negative correlation (ρ<-0.7) for CYPs, GTS, NHRs in *H. bakeri* against the duplication rate (Table S44). The other comparisons could not be considered noteworthy (|ρ|<0.7). Across the 628 clusters that we generated, approximately 60% contained the same number of members for each assembly. Put another way, ∼40% of clusters contained a variable number of proteins and often this variation was an additional member in just one assembly (Tables 6 and S45-S68).

We provide two examples of variable cluster memberships from the CYPs. The first is cluster bovis-cyp-0000003 (Table S45 and Table S46), in which: *C. elegans* and the Redbean2.5 assembly have one member each; the Stevens, Canu and Flye assemblies have two members each; the Falcon and Falcon-unzip assemblies have 3 members each. The *C. elegans* protein is CYP-33C9 whose function has been associated with fatty-acid desaturation hence reducing susceptibility of xenobiotics by increasing desiccation tolerance (Erkut et al., 2013). The second example is cluster contortus-cyp-0000026, in which: *C. elegans*, in which: *C. elegans* and the Flye, Falcon, and Falcon-unzip assemblies have one member each; the Doyle, Canu, Redbean2.5, and SMARTdenovo assemblies have no members (Table S47 and Table S48). The *C. elegans* protein is CYP-43A1 and is highly expressed in the intestine and serotonergic neuron ADF (Packer et al., 2019). Interestingly, an orthologue to C. elegans CYP-43A1 was found in an earlier version of the *H. contortus* assembly and shown to be the most highly expressed CYP in the *H. contortus* intestine (Laing et al., 2015).

In *H. bakeri*, three protein families—TGF-b mimics (TGM), ARI, and BARI—have been shown to modulate the host’s immune system (Osbourn et al., 2017; Johnston et al., 2017; Smyth et al., 2018). The available protein sequences for these families aligned poorly to the *H. bakeri* Braker3 gene models (data not shown). We determine the likely locations of these proteins on the genome assemblies using two splice-aware aligners, exonerate and miniprot (Slater & Birney, 2005; Li, 2023). Similar to the other protein families, we found variation in gene counts across the three HiFi long-read assemblies: nine-to-16 TGMs and 13-to-23 ARIs/BARIs (Table 7 and Table S69). We checked whether the protein-to-genome alignments overlapped with Braker3 gene models and found that while most alignments (TGM 81-100%; ARI/BARI 68-100%) overlapped with a gene model, far fewer alignments (TGM 40-91%; ARI/BARI 27-67%) overlapped gene models across by greater than 50% of their length (Table 7). Focusing on scaffold ptg000126l from the Hifiasm assembly, we saw a full set of examples, from complete congruence between Braker3, exonerate, and miniprot, to evidence from just one source (Figure 6).

**Figure 6:**
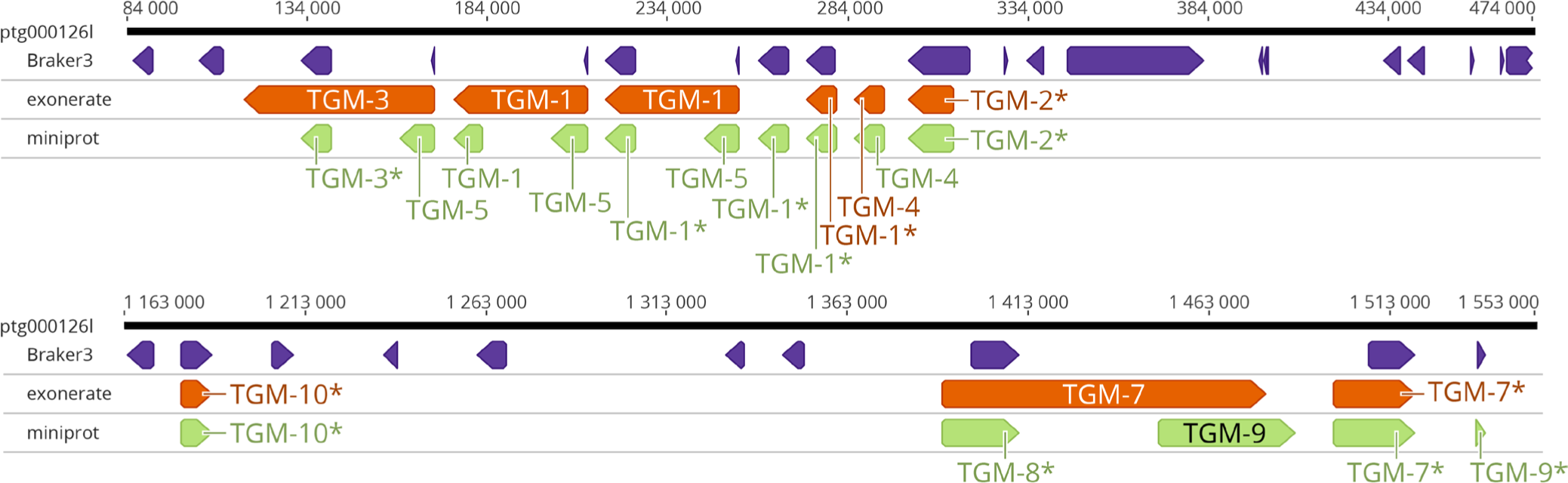
Location of TGM protein-coding genes on *H. bakeri* scaffold ptg000126l. TGM genes were found to cluster in two locations on this scaffold. The purple track is the Braker3 gene model predictions, the orange track is the Exonerate protein-to-genome alignments, the green track is the miniprot protein-to-genome alignments. For both exonerate and miniprot alignments, an * signifies ≥50% reciprocal overlap with a Braker3 gene model. The black bar is the scaffold, with nucleotide coordinates given above the bar.

**Table 7:** Immunomodulators annotated in *H. bakeri* assemblies.

| Assembly | Chow |  | Flye |  | HiCanu |  | Hifiasm |  |
| --- | --- | --- | --- | --- | --- | --- | --- | --- |
| Algorithm | Miniprot | Exonerate | Miniprot | Exonerate | Miniprot | Exonerate | Miniprot | Exonerate |
| TGM-1 | 0 | 0 | 1 | 1 | 2 | 1 | 4 | 2 |
| TGM-2 | 1 | 1 | 2 | 1 | 1 | 1 | 1 | 1 |
| TGM-3 | 0 | 0 | 0 | 0 | 1 | 1 | 1 | 1 |
| TGM-4 | 1 | 1 | 1 | 1 | 1 | 1 | 1 | 1 |
| TGM-5 | 1 | 1 | 1 | 1 | 1 | 1 | 3 | 1 |
| TGM-6 | 1 | 1 | 1 | 1 | 1 | 1 | 1 | 1 |
| TGM-7 | 1 | 1 | 1 | 1 | 1 | 2 | 1 | 2 |
| TGM-8 | 1 | 1 | 2 | 1 | 1 | 0 | 2 | 0 |
| TGM-9 | 2 | 2 | 1 | 1 | 1 | 1 | 1 | 0 |
| TGM-10 | 1 | 1 | 1 | 1 | 1 | 1 | 1 | 1 |
| **Total TGMs** | **9** | **9** | **11** | **9** | **11** | **10** | **16** | **10** |
| **Overlap gene model (any)** | **9** | **9** | **11** | **9** | **11** | **10** | **13** | **9** |
| **Overlap gene model (≥50%)** | **7** | **6** | **10** | **6** | **10** | **8** | **9** | **4** |
| ARI1 | 1 | 1 | 3 | 2 | 7 | 3 | 8 | 4 |
| ARI2 | 1 | 1 | 1 | 2 | 1 | 2 | 1 | 2 |
| ARI3 | 4 | 3 | 3 | 3 | 3 | 3 | 3 | 3 |
| BARI | 2 | 1 | 2 | 2 | 3 | 3 | 3 | 3 |
| BARI_Hom1 | 2 | 4 | 3 | 5 | 7 | 4 | 7 | 4 |
| BARI_Hom2 | 1 | 1 | 1 | 1 | 1 | 1 | 1 | 1 |
| **Total ARI/BARI** | **11** | **11** | **13** | **15** | **22** | **16** | **23** | **17** |
| **Overlap gene model (any)** | **10** | **10** | **13** | **14** | **15** | **14** | **17** | **14** |
| **Overlap gene model (≥50%)** | **3** | **3** | **6** | **10** | **10** | **8** | **8** | **8** |

## Discussion

Improvements in sequencing read length and accuracy is fuelling the demand for more contiguous genomes and led to the development of many novel computational strategies. The technologies are rapidly evolving, and we have not atempted to be comprehensive; rather, we selected genome assemblers based on their use at the start of our study. Benchmarking in other organisms has shown that there remains value in comparative analyses, especially given the remarkable genetic diversity encountered across the genomes sequenced to date (Pollo et al., 2020; Jung et al., 2020; Sun et al., 2021; Koo et al., 2023). Our objectives here have been to determine the relative efficacy of different assembly programs, particularly their effect on gene identification within nematodes species. These species cover a wide range of genomic characteristics, which should make our findings of interest to those sequencing and assembling their own favourite, non-nematode, organisms.

Long-read assemblers predominantly use two core algorithmic frameworks: Overlap-Layout-Consensus (OLC) and De-Bruijn Graph (DBG) (reviewed in (Li et al., 2012)). Each framework is decorated with a myriad of developer adaptations to address specific genome features or challenges, such as repetitive sequences, varying levels of ploidy, and differential sequencing error profiles. The process of implementing and optimising these adaptations is not trivial; they significantly affect the assemblers’ effectiveness, especially in regions exhibiting intra-population structural complexities or high heterogeneity.

Arguably, the least complex genome in this study was *C. bovis* and DBG-based assemblers—Redbean2.5 and Flye—performed best, yielding less fragmented assemblies and higher BUSCO scores. They also did beter with the more granular measures of assembly performance: read mapping rates, alignment depths, and structural errors (large and small). Determining the best assemblies for *H. contortus* was less obvious. Here, OLC-based assemblers—SMARTdenovo and Canu—achieved larger N50s and marginally beter BUSCO scores. The Redbean2.5 (DBG) assembly had the lowest BUSCO duplication rate and fewest structural errors. For *H. bakeri*, the Hifiasm assembler, produced the most contiguous assembly but according to the Inspector report displayed the highest number of structural errors. This suggests potential challenges in handling highly heterozygous regions by the assembly algorithm, despite its default haplotype-resolving capabilities.

From our reading of the literature, the BUSCO score may be the most widely used metric to determine assembly accuracy, and we too relied heavily on it in the presented analysis. Often, we found that the scores were largely similar between assemblers, suggesting that the gene complements of each genome would be similar. However, diving deeper into the classification of the 3131 nematode single copy orthologous demonstrated a surprising situation, where a gene may be found as a single copy in half the assemblies and missing in another half. We saw differences between Redbean versions 2.3 and 2.5 which should motivate researchers to carefully consider updated annotations; that is to ask, what gets lost when you improve your annotation? We found at least 27 genes which were potentially incorrectly absent in the published *H. contortus* assembly. This is one of the most carefully curated parasite genome assemblies and demonstrated the need to fund ongoing curation for important species (Doyle et al., 2020; Doyle, 2022).

The reliance on BUSCO scores is that small variations in score can be considered trivial. For example, 1% change in nematodes is 31 genes. Does this mater? It is important to remember that BUSCO genes are the most conserved genes in an organism. They typically perform critical functions and their absence from an assembly, if true *in vivo*, would point to a significant change in an organism’s biology. However, If the absence is due to a technical problem correctly assembling the reads, one might assume that assembly of regions containing less well conserved single copy genes would be more error-prone, particularly for multi-copy gene families. This is what we find with the *cyp*, *gst*, *ugt*, and *nhr* gene family. Comparative analyses often involve species assembled and annotated using different methods across different laboratories (Coghlan et al., 2019). This annotation heterogeneity inflates the number of lineages-specific genes (Weisman, Murray & Eddy, 2022). An alternative is to reannotate the assemblies with a uniform approach, as we did here. A limitation of this approach, and our study, is that we then ignore expert curation. For example, our reannotation of the Doyle *H. contortus* assembly resulted in approximately 4,800 genes then the published annotation, and a lower BUSCO score. In contrast, our reannotation of the published *C. bovis* and *H. bakeri* assemblies resulted in more genes, and importantly, improved BUSCO scores. Together, this demonstrates the requirement to validate the absence of a gene, which may actually be present in the assembly but score just the wrong side of a gene finder’s score cut-off (Gilabert et al., 2016; Stroehlein, Young & Gasser, 2018; Baker et al., 2021).

The TGM proteins were discovered through a proteomic survey of *H. bakeri* secreted products and later mapped to the Chow assembly, where only two of the ten proteins could be confidently assigned to gene models and another two proteins mapped directly to the genome (Johnston et al., 2017; Smyth et al., 2018). Similarly, the alarmin release inhibitors (ARI and BARI) evaded direct mapping to the published Chow assembly gene models. We find that even with improved assemblies, these proteins could not be reliably reconstructed with arguably the most popular gene finding software. However, targeted searches did provide strong evidence for genes for all proteins, some with multiple gene copies. However, as the variation across the assemblies shows, a more careful curation is required.

Determining the usefulness of an assembly will remain a personal choice; one that should be guided by a range of metrics. It is noteworthy that in a comparison of assemblies generated for several invertebrates, the assemblies with the highest BUSCO score were sometimes the most fragmented (Sutton et al., 2021).

## Conclusion

We encourage those charged with assembling genomes—be it *Heligmosomoides*, *Haemaphysalis*, or *Henneguya*—to be aware of the important differences between seemingly very similar assemblies. For users of genome assemblies and annotation, we emphasize the importance of exploring alternative assemblies generated by multiple assemblers to validate biological hypotheses. Finally, we note that much of the above discussion can be summed up neatly by the first recommendation from Assemblethon2: “Don’t trust the results of a single assembly (Bradnam et al., 2013).”

## Acknowledgements

We acknowledge the high-performance computing resources made available by the Faculty of Veterinary Medicine and Research Computing at the University of Calgary. We thank Drs. Stephen Doyle, John Gilleard, Graham Plastow, John Soghigian, and Frank van der Meer for discussions around this topic.

## Author contribution

GMM: Conceptualization; Data curation; Formal analysis; Investigation; Methodology; Validation; Visualization; Writing – original draft; Writing – review & editing

JDW: Conceptualization; Formal analysis; Funding acquisition; Investigation; Methodology; Supervision; Validation; Visualization; Writing – original draft; Writing – review & editing

## Data availability

Sequence reads for the *Heligmosomoides bakeri* genome are available on the sequence read archive (SRA): SRX23938287 and SRX23938287. File S2 contains the genome assemblies and Braker3 gene models generated in this study and is available on Datadryad (doi.org/10.5061/dryad.p2ngf1vzh)

