## Supplementary figures and images for "Genome assembly variation and its implications for gene discovery in nematode species"

### Figure S1

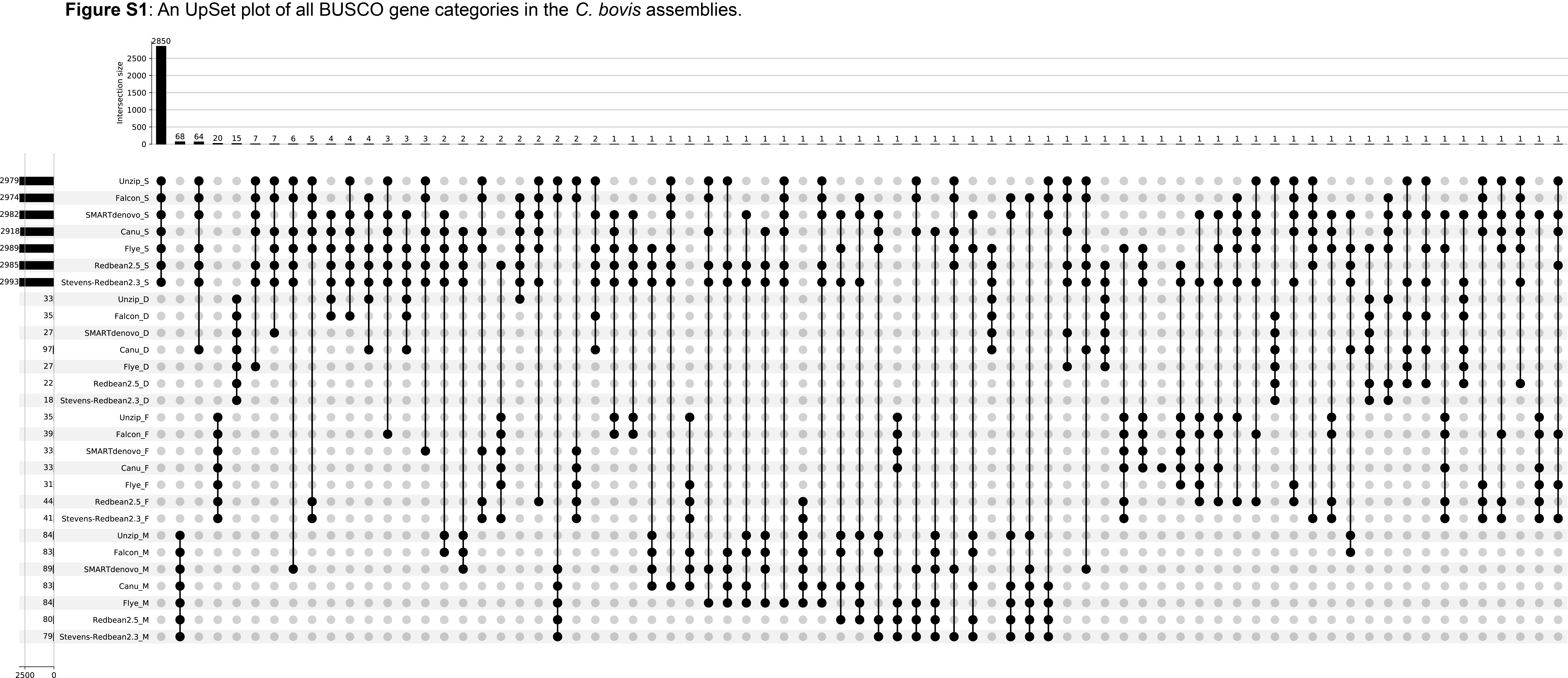

### Figure S2

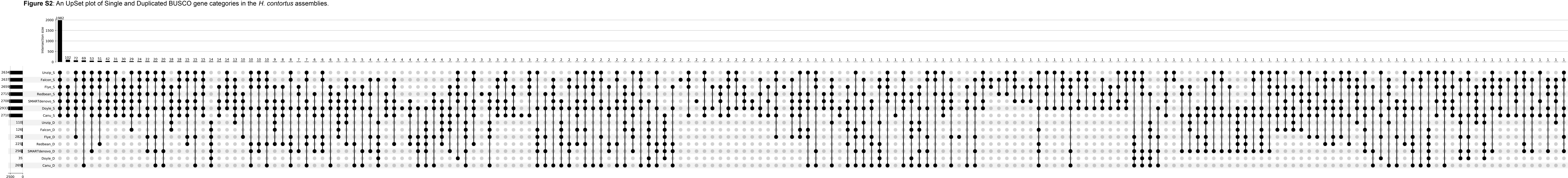

### Figure S3

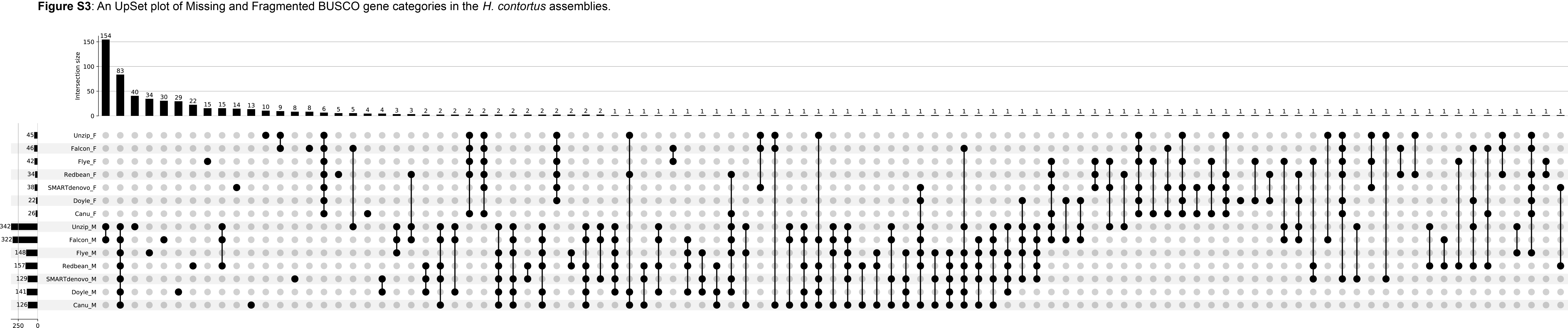

### Figure S4

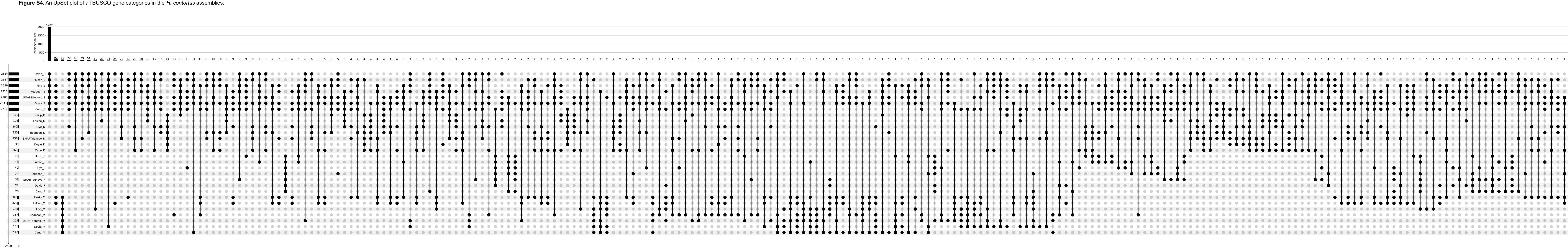

### Figure S5

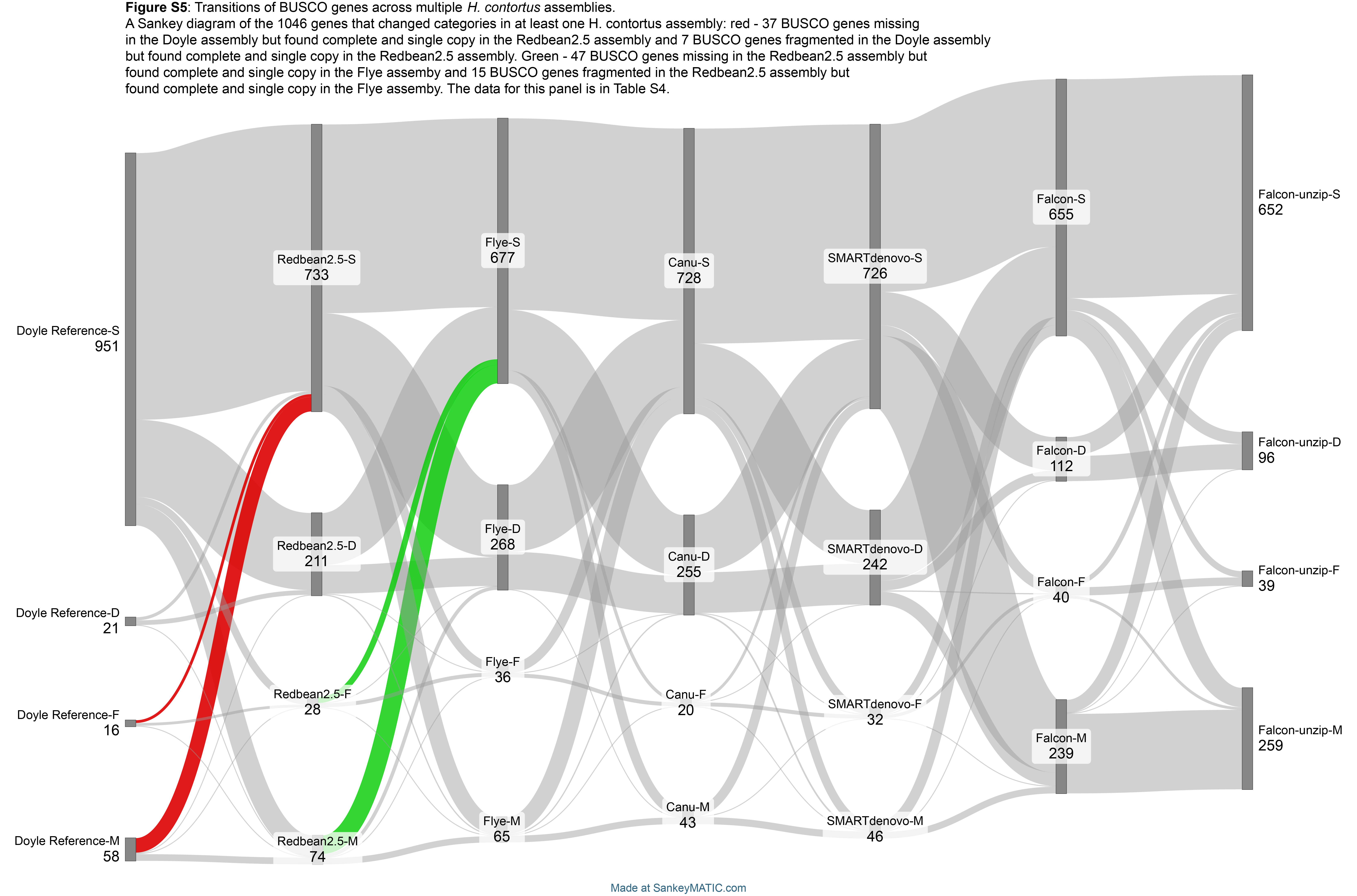

### Figure S6

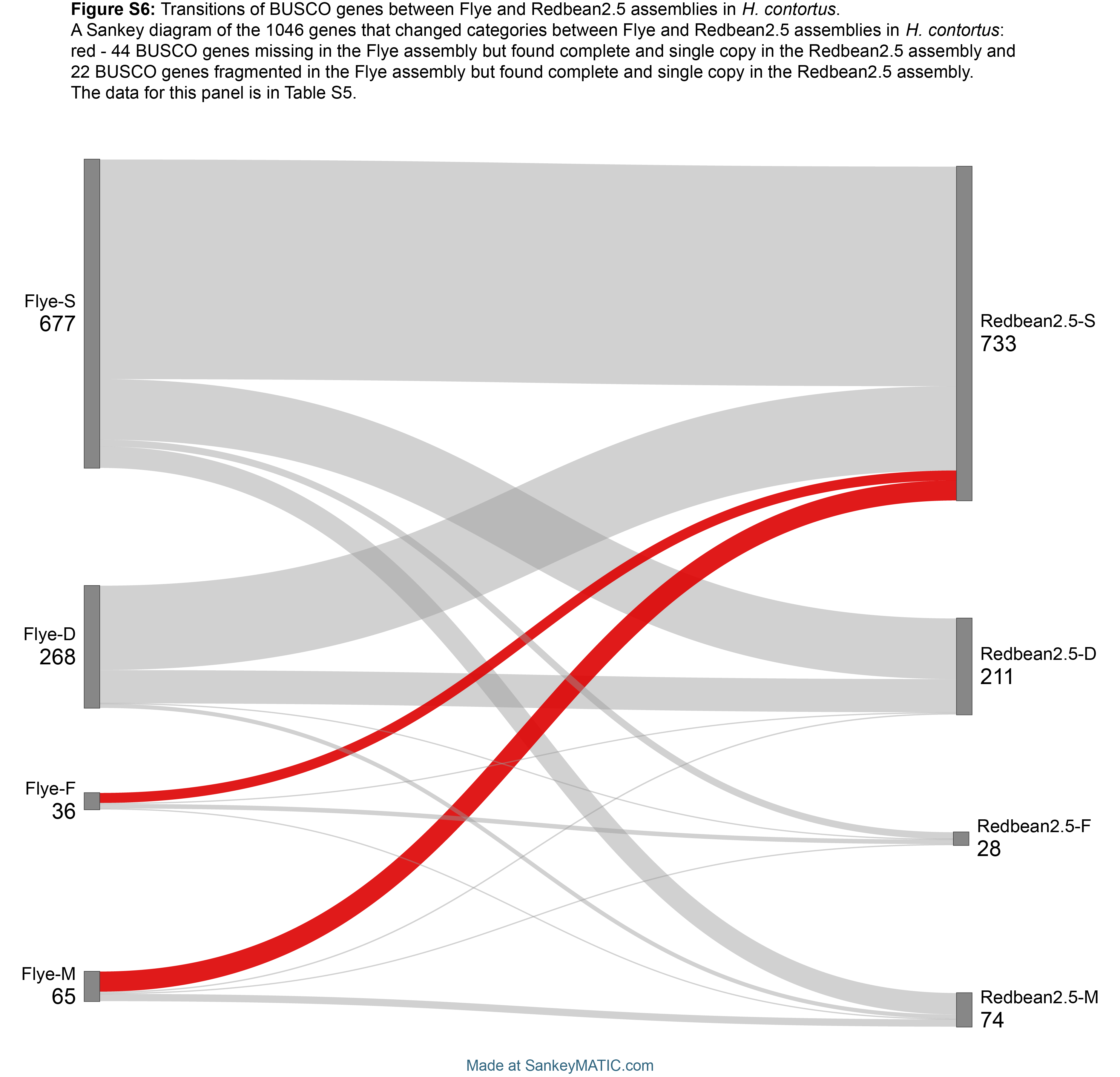
