## Supplementary material for "Genome assembly variation and its implications for gene discovery in nematode species": File S1

**GENOME ASSEMBLY COMMANDS**

```
## Commands used, replace anything in square brackets [] with your appropriate path or unique name
## Commands to assemble preliminary long read assemblies
## C.bovis and H.contortus assemblies
## Canu:
# For C.bovis:
canu -p [UNIQUE_NAME] -d [UNIQUE_NAME] genomeSize=62.7m useGrid=true gridOptions="--time=24:00:00" -nanopore [PATH/TO/READS]
# For H.contortus:
canu -p [UNIQUE_NAME] -d [UNIQUE_NAME] genomeSize=283.4 m useGrid=true gridOptions="--time=24:00:00" -pacbio [PATH/TO/READS]

## Flye:
# For C.bovis:
flye --nano-raw [PATH/TO/READS] --out-dir [UNIQUE_NAME] --threads [12]
# For H.contortus
flye --pacbio-raw [PATH/TO/READS] --out-dir [UNIQUE_NAME] --threads [12]

## Redbean (also known as wtdbg2):
# For C.bovis:
wtdbg2 -x ont -g 62.7m -t [8] -i [PATH/TO/READS] -o [UNIQUE_NAME]
wtpoa-cns -t [8] -i [UNIQUE_NAME].ctg.lay.gz -o [UNIQUE_NAME]
# For H.contortus:
wtdbg2 -x ont -g 283.4m -t [8] -i [PATH/TO/READS] -o [UNIQUE_NAME]
wtpoa-cns -t [8] -i [UNIQUE_NAME].ctg.lay.gz -o
[UNIQUE_NAME]

## SMARTdenovo:
smartdenovo.pl -p [UNIQUE_NAME] -c 1 [PATH/TO/READS] > [UNIQUE_NAME].mak
make -f [UNIQUE_NAME].mak
wtcns -t 16 < [UNIQUE_NAME].dmo.lay > [UNIQUE_NAME].fasta

## Falcon:
## Modify the default configuration file according to compute environment and input file:
      #### Input
      [General]
      input_fofn=input.fofn ## Your list of paths to the input fasta files
      input_type=raw
      pa_DBdust_option= ## By default, dusting is turned on
      pa_fasta_filter_option=streamed-median
      target=assembly
      skip_checks=False
      LA4Falcon_preload=false
      #### Data Partitioning
      ### Large genomes(>10Mb)
      pa_DBsplit_option=-x500 -s200
      ovlp_DBsplit_option=-s200
```

```
    #### Repeat Masking
    pa_HPCTANmask_option=
    #no-op repmask param set
    pa_REPmask_code=0,300;0,300;0,300
    ####Pre-assembly
    ## adjust to your genome size
    genome_size = 62726283 ## for C.bovis
    ## OR genomesize = 283439308 ## for H.contortus
    seed_coverage = 40
    length_cutoff = -1
    pa_HPCdaligner_option=-v -B128 -M24
    pa_daligner_option= -k18 -e0.80 -l3000 -h256 -w8 -s100
    falcon_sense_option=--output-multi --min-idt 0.70 --min-cov 4 --max-
         n-read 400
    falcon_sense_greedy=False
    ####Pread overlapping
    ovlp_HPCdaligner_option=-v -B128 -M24
    ovlp_daligner_option= -k24 -e.94 -l3000 -h1024 -s100
    ####Final Assembly
    length_cutoff_pr=1000
    overlap_filtering_setting=--max-diff 100 --max-cov 100 --min-cov 2
    fc_ovlp_to_graph_option=
    [job.defaults]
    job_type=slurm
    pwatcher_type=blocking
    JOB_QUEUE=[settings for your default job queue/partition]
    MB=300 ## memory allocated per job
    NPROC=16 ## number of processors per job
    njobs=10 ## number of concurrently running jobs
    submit = srun --wait=0 -p {JOB_QUEUE} \ ## for slurm use srun
    -J ${JOB_NAME}        \
    -o ${JOB_STDOUT}      \
    -e ${JOB_STDERR}        \
    --mem-per-cpu=${MB}M \
    --ntasks 1   \
    --cpus-per-task=${NPROC} \
    ${JOB_SCRIPT}
## Running Falcon
##create file of file names
readlink -f [READS] > input.fofn
fc_run [UNIQUE_CONGIG_NAME].cfg
```

**## Falcon-unzip:**
**##Falcon-unzip config file:**

```
    [General]
     max_n_open_files = 500
    [Unzip]
    input_fofn=input.fofn
    input_bam_fofn=input_bam.fofn
    polish_include_zmw_all_subreads = true
```

```
      [job.defaults]
      job_type=slurm
      pwatcher_type=blocking
      JOB_QUEUE=[settings for your default job queue/partition]
      MB=300
      NPROC=4
      njobs=8
      submit = srun --wait=0 -p ${JOB_QUEUE}  \
      -J ${JOB_NAME}        \
      -o ${JOB_STDOUT}       \
      -e ${JOB_STDERR}        \
      --mem-per-cpu=${MB}M     \
      --cpus-per-task=${NPROC} \
      ${JOB_SCRIPT}
##Running Falcon-unzip
readlink -f *.fasta > input.fofn
readlink -f *.bam > input_bam.fofn
fc_unzip.py [UNIQUE_CONGIG_NAME].cfg
```

**##Preliminary assemblies for *H.bakeri***

**## Hicanu**

```
canu -assemble -p [UNIQUE_NAME]-d [UNIQUE_NAME] genomeSize=696.9m
useGrid=true cnsMemory=64 batMemory=62 gridOptions="--time=7-00:00:00" -
pacbio-hifi [PATH/TO/READS]
```

**## Hifiasm**

```
hifiasm -o [UNIQUE_NAME] -t 24 [PATH/TO/READS]
```

**## Flye**

```
flye --pacbio-hifi [PATH/TO/READS] --out-dir [UNIQUE_NAME] --threads [12]
```

**DECONTAMINATION AND POLISHING ASSEMBLIES**

**## Mapping of the long reads to assemblies using Minimap2**

```
minimap2 -ax map-ont \ ## use -ax map-pb for H.contortus
[PATH/TO/ASSEMBLY] [PATH/TO/READS] > [UNIQUE_NAME].sam
# Sorting the alignments
samtools view -b [UNIQUE_NAME].sam | sort > [UNIQUE_NAME].sorted.bam
samtools index [UNIQUE_NAME].sorted.bam
```

**## Running Diamond**

**# Extract and concatenate taxid mapping files from NCBI**

```
echo "accession\taccession.version\ttaxid\tgi" >
reference_proteomes.taxid_map
zcat */*.idmapping.gz | grep "NCBI_TaxID" | awk '{print $1 "\t" $1 "\t" $3
"\t" 0}' >> reference_proteomes.taxid_map

wget -N ftp://ftp.ncbi.nlm.nih.gov/pub/taxonomy/taxdump.tar.gz
```

```
mkdir -p taxdump && tar zxf taxdump.tar.gz -C ./taxdump

# Make diamond blast database with taxonomic information
diamond makedb --in [concatenated-reference_proteomes.fasta.gz] --taxonmap reference_proteomes.taxid_map \
 --taxonnodes [PATH/TO/taxdump/nodes.dmp] --db reference_proteomes.dmnd

diamond blastx --query [PATH/TO/ASSEMBLY] \
 --db [PATH/TO/reference_proteomes.dmnd] \
 --outfmt 6 qseqid staxids bitscore qseqid sseqid pident length mismatch gapopen qstart qend sstart send evalue bitscore \
 --max-target-seqs 1 --evalue 1e-25 -o [UNIQUE_NAME].tsv --very-sensitive

## Running BlobTools
blobtools map2cov -i [PATH/TO/ASSEMBLY] \
 --bam [PATH/TO/MINIMAP2_SORTED_BAM_FILE] -o [UNIQUE_NAME_FOR_COVERAGE_FILE]

blobtools create -i [PATH/TO/ASSEMBLY] -t [PATH/TO/DIAMOND_OUTPUT_FILE.tsv] \
 --db [PATH/TO/BLOBTOOLS/nodesDB.txt] -x bestsumorder \
 --cov [UNIQUE_NAME_FOR_COVERAGE_FILE] -o [UNIQUE_NAME]

## Mapping and polishing decontaminated assemblies
# Aligning the long reads to the filtered assembly using minimap2 and performing four rounds of racon.
minimap2 -ax map-ont \ ## use -ax map-pb for H.contortus
[PATH/TO/ASSEMBLY] [PATH/TO/READS] > [UNIQUE_NAME].sam

racon -m 8 -x -6 -g -8 -w 500 -t 16 [PATH/TO/READS] \
[PATH/TO/MINIMAP_OUTPUT.sam] \
[PATH/TO/BLOBTOOLS_OUPUT.fasta]
> [UNIQUE_NAME_FOR_RACON_ROUNDS].fasta
## Repeat these steps four times with the output from the first round as input for the second round and so on. After the fourth round then:

## For C. bovis: Polishing racon_round4.fasta using Medaka.
medaka_consensus -i [PATH/TO/READS] \
 -d [PATH/TO/RACON_ROUND4.fasta] \
 -o [UNIQUE_NAME_FOR_MEDAKA_CONSENSUS] -t 24 -m r941_min_high_g303

## For H. contortus: Polishing racon_round4.fasta using Arrow.
## First map using pbmm2

pbmm2 align [PATH/TO/RACON_ROUND4.fasta] \
[PATH/TO/HCONTORTUS_SUBREADS.bam] \
[UNIQUE_OUTPUT_NAME].bam --sort -j 12 -J 4

pbindex [UNIQUE_OUTPUT_NAME].bam
```

**## Running Arrow**
```
samtools faidx [PATH/TO/RACON_ROUND4.fasta]

arrow -j 32 [PATH/TO/PBMM2_OUTPUT.bam] -r [PATH/TO/RACON_ROUND4.fasta] \
 -o [UNIQUE_OUTPUT_NAME].gff -o [UNIQUE_OUTPUT_NAME].fasta
```

**## Mapping Illumina short reads to the Medaka/Arrow polished assemblies using BWA-MEM and performing two iterations of racon.**
**## For *C.bovis*:** bwa index [UNIQUE_NAME_FOR_MEDAKA_CONSENSUS].fasta
**## For *H.contortus*:** bwa index [UNIQUE_NAME_FOR_ARROW_OUPUT].fasta

bwa mem -t 24 [UNIQUE_NAME_FOR_MEDAKA_CONSENSUS].fasta **## Or for *H.contortus*:[UNIQUE_NAME_FOR_ARROW_OUPUT].fasta**
[PATH/TO/ILLUMINA/SHORT/READS] > [UNIQUE_NAME].sam

**racon** -m 8 -x -6 -g -8 -w 500 -t 24 \
[PATH/TO/ILLUMINA/SHORT/READS] \
[PATH/TO/BWA-MEM_OUTPUT].sam \
[UNIQUE_NAME_FOR_MEDAKA_CONSENSUS].fasta \ **## Or for *H.contortus*:[UNIQUE_NAME_FOR_ARROW_OUPUT].fasta**
> [UNIQUE_NAME_FOR_RACON_ROUNDS].fasta

**## Repeat these steps two times with the output from the first round as input for the second round. After the second round then:**
**## Performing two iterations of pilon on the racon_round2.fasta assembly.**

```
bwa index [PATH/TO/RACON_ROUND2.fasta]

bwa mem -t 8 [PATH/TO/RACON_ROUND2.fasta] \
[PATH/TO/ILLUMINA_READ1] [PATH/TO/ILLUMINA_READ2 | samtools sort -O bam \
-o [UNIQUE_NAME_FOR_SORTED].bam -T [UNIQUE_NAME_FOR_TEMP_FILE]

samtools index [UNIQUE_NAME_FOR_SORTED].bam


java -Xmx60G -jar pilon-1.24.jar \
 --genome [PATH/TO/RACON_ROUND2.fasta] --fix bases --frags
[PATH/TO/SORTED/BAM/FILE]
--threads 24 --output [UNIQUE_NAME_PILON_ROUNDS] | tee
[UNIQUE_NAME_PILON_ROUNDS]
```

**## Repeat these steps two times with the output from the first round as input for the second round.**

**HAPLOTYPE DUPLICATION REMOVAL**
**## Purge_dups commands and parameters**
```
pri_asm=[PATH/TO/PRIMARY/ASSEMBLY].fasta
pb_fasta=[PATH/TO/READS]
```

```
ref_genome_split=[PATH/TO/SPLIT/PRIMARY/ASSEMBLY].split.fasta
self_aln_genome=[PATH/TO/SELF/ALIGNED/SPLIT/PRIMARY/ASSEMBLY].split.self.paf.gz
```

**## Runnng minimap2 to align pacbio data and generating paf files, then calculating read depth histogram and base-level read depth**

```
minimap2 -I 6G -x map-pb $pri_asm $pb_fasta | gzip -c - > purge.paf.gz
```

**## Producing PB.base.cov and PB.stat files**

```
[PATH/TO/BIN/pbcstat] [ALL].paf.gz
[PATH/TO/BIN/CALCUTS PB.stat] > cutoffs 2> calcults.log
python [PATH/TO/SCRIPTS/hist_plot.py -c cutoffs PB.stat PB.png
```

**## Splitting the primary assembly and doing a self-self alignment**

```
[PATH/TO/BIN/split_fa $pri_asm > $pri_asm.split.fa
[PATH/TO/BIN/split_fa $pri_asm > $ref_genome_split

minimap2 -x asm5 -DP $ref_genome_split $ref_genome_split | gzip -c - > $self_aln_genome
```

**## Purging haplotigs and overlaps**

```
[PATH/TO/BIN/purge_dups -2 -T cutoffs -c PB.base.cov $self_aln_genome > dups.bed 2> purge_dups.log
```

**## Getting purged primary and haplotig sequences from draft assembly**

```
[PATH/TO/BIN/get_seqs dups.bed $pri_asm > [PURGED_ASSEMBLY].fa 2> [HAPLOTIG_SEQUENCES].fa
```

**ASSEMBLY EVALUATIONS**

**## BUSCO command used to evaluate the completeness of single copy orthologues:**

```
busco -i [PATH/TO/ASSEMBLY] --lineage nematoda_odb10  --out [UNIQUE_NAME]  --mode genome --long –augustus
```

**## Additionally, we assessed the alignment of sequence reads to each assembled genome using Inspector**
**## For this evaluation, we used the raw ONT reads for the *C. bovis* assemblies, the raw PacBio RS and Sequel reads for the *H. contortus* assemblies, and the raw HiFi long reads for the *H. bakeri* assemblies.**
**## Inspector commands used to align sequence reads to assemblies to identify both large- and small-scale errors.**

inspector.py --thread 24 --contig [PATH/TO/ASSEMBLY] --read [PATH/TO/READS] --outpath [UNIQUE_NAME_FOR_OUTPUT_DIRECTORY] --datatype [nanopore for *C. bovis* or clr for *H. contortus* or hifi for *H. bakeri*].

**## To generate the corrected assembly**

```
inspector-correct.py --thread 24 --inspector [UNIQUE_NAME_FOR_OUTPUT_DIRECTORY] --datatype [nano-raw or pacbio-raw or pacbio-hifi] --outpath [DIRECTORY_FOR_CORRECTED_ASSEMBLY].
```

**GENOME SYNTENY**

```
## Genome-to-genome alignments of the final draft assemblies using the Nucmer
nucmer --maxmatch --prefix=[UNIQUE_NAME] [REFERENCE GENOME] [QUERY GENOME]

## For the C. bovis, we used the dnadiff wrapper script to report one-to-one alignment coordinates
dnadiff -d [NUCMER_OUTPUT].delta

## For all the generated assemblies, we used NucDiff to quantify the structural and local genome differences between each assembly and its respective reference genome
nucdiff --proc 24 --ref_name_full yes --query_name_full yes [REFERENCE GENOME] [QUERY GENOME] [PREFIX NAME]
```

**GENE PREDICTION**

```
##RNA-seq data alignment using STAR (For H. contortus and H. bakeri)
STAR --runThreadN [24] --runMode genomeGenerate \
 --genomeDir [DIRECTORY_FOR_ASSEMBLY_INDEX]
 --genomeFastaFiles [PATH/TO/ASSEMBLY]

## Mapping RNA-seq reads
STAR --genomeDir [ASSEMBLY_INDEX_DIRECTORY] --readFilesCommand zcat \
 --readFilesIn [INPUT RNA-SEQ READS] --runThreadN [20] \
 --outSAMtype BAM Unsorted \
 --outFileNamePrefix [UNIQUE_NAME]

## Sorting the aligned RNA-seq bam file
samtools sort [UNIQUE_NAME.bam] --threads [20] -o [UNIQUE_NAME_sorted.bam] -T tmp.bam
```

**BRAKER 3 RUN**

```
export BRAKER_SIF=[PATH/TO/SINGULARITY/CONTAINER]
export GENEMARK_PATH=[PATH/TO/GENEMARK]
export PATH=[PATH/TO/PROTHINT/BIN]:$PATH

singularity exec -B $PWD:$PWD [PATH/TO/SINGULARITY/CONTAINER] \
cp -r /usr/share/augustus/config $PWD/.augustus \
singularity exec -B $PWD:$PWD [PATH/TO/SINGULARITY/CONTAINER] \
braker.pl --AUGUSTUS_CONFIG_PATH=$PWD/.augustus  \
 --species=[SPECIES_NAME] --threads=24 --softmasking \
 --genome=[PATH/TO/SOFTMASKED/GENOME] \
--GENEMARK_PATH=[PATH/TO/GENEMARK]
 --bam=[PATH/TO/SORTED/RNA-SEQ.bam] ## For H. contortus and H. bakeri
 --prot_seq=[PATH/TO/PROTEIN/SEQS.fasta] ## For C. bovis
--PROTHINT_PATH=[PATH/TO/PROTHINT/BIN] ## For C. bovis
```
